# Auxin regulates the proportions, but not the initiation, of *Hibiscus trionum* bullseyes

**DOI:** 10.64898/2026.11.19.767829

**Authors:** Elena Salvi, Stefano Gatti, May T. Yeo, Lucie Riglet, Erin Doody, Edwige Moyroud

## Abstract

Colourful motifs on the surface of flower petals fulfil important ecological functions, mediating pollination and providing protection against environmental stressors. These patterns form when epidermis cells in different subdomains of the petal acquire contrasting chemical and structural features. While some of the enzymes and transcription factors involved in specifying the production of different pigments, cell shapes and textures have been identified, the signalling events and upstream processes that divide the petal surface in distinct regions are not yet understood. Plant hormones play key roles in orchestrating all aspects of plant development but whether and how they contribute to the establishment of petal patterns remain to be tested. Here, we investigate the roles of auxin during bullseye pattern development in *Hibiscus trionum*. We found that auxin does not initiate bullseye boundary formation during the pre-patterning phase. Instead, auxin signalling is essential for regulating cell behaviour in the distal region and preserving the bullseye proportions established in young petal primordia. Therefore, auxin could act as a pattern modifier over evolutionary timescales, shifting bullseye dimensions by modulating the relative growth of different petal regions.

## Introduction

Angiosperms can generate a huge array of petal epidermal patterns and shapes that serve a multitude of vital functions. Petal patterns, the colourful motifs that ornate the corolla of flowering plants, are macroscopically visible for animals and humans mainly as pigmentation, texture and iridescent patterns ^1,2^. They can act as aposematic signals to repel herbivores and as advertising cues to lure pollinators ^3^. UV-absorbing patterns can also protect the flower reproductive organs and cells against damaging radiations ^4,5^ and desiccation ^6^.

Petal patterns are generated and organised at the cellular and histological scale due to differential pigment accumulation in cell organelles, to different cell shapes and to cuticular ornamentations of the cells across the petal epidermis ^7,8^. Petal patterns also result from differences in space and time of cell activities (cell proliferation, cell expansion and cell differentiation), with growth acting as a modifier of (pre-)patterns set early during development ^9–11^. Hence, alterations in petal patterns, happening at any stage of petal development, could yield direct consequences for the plant ability to reproduce and its fitness, thereby causing potentially severe consequences in agriculture, economy and society. Nonetheless, how patterns are generated across the surfaces of living organisms is still not fully understood ^12,13^.

Small mobile molecules are key regulators of morphogenesis in multicellular organisms, initiating organ primordia, establishing developmental axes, regulating spatio-temporal growth and specifying cell types ^14,15^. In animals, secreted factors can act as mitogens, morphogens and cell differentiation cues according to their local concentration gradients, timing and the molecular context: for instance, Sonic Hedgehog patterns the embryonic ventral neural tube, the posterior limb bud, the embryonic somites, foregut, lung and brain, and regulates the craniofacial development in vertebrates ^16^. Similarly, phytohormones shape endogenous developmental programmes, and coordinate them by mediating tropisms in response to exogenous cues ^17,18^. Plant hormones are central in initiating, shaping and patterning plant tissues and organs although clear and specific evidence is still lacking regarding the roles phytohormones play during petal patterns formation ^10^.

Pattern formation requires the establishment of axes and developmental boundaries on which pattern outlines are laid accordingly ^19,20^. Auxin regulates virtually all plant developmental events and as such it is the most extensively studied phytohormone. Multiple experimental tools like auxin analogues, inhibitors of specific signal transduction components, and reporters for *in vivo* activity tracking have been developed ^21,22^. Auxin is distributed differentially in tissues, and its homeostasis, transport and signalling machinery are finely spatio-temporally regulated at the transcriptional, post-transcriptional and post-translational levels ^23–25^. Auxin maxima, minima and gradients act as morphogenetic triggers and positional information signals in main and lateral organs ^26^, like in the shoot apical meristem to distinguish cell fates within the same apical organ ^27^ or to specify the founder cell for lateral root initiation ^28,29^. In the root apical meristem, an auxin minimum separates different cell behaviour longitudinally ^30^. Auxin distribution set developmental boundaries, including those between floral whorls and within floral whorls ^31,32^ or during embryogenesis, between apical meristems and emerging cotyledons ^33,34^. Remarkably, auxin is involved in the control of cell growth by controlling cell-cycle progression and cell division rates ^35–38^, cell division patterns and orientations in Arabidopsis embryos and roots ^39,40^, cell expansion in leaves, roots and hypocotyls ^41–43^and growth axes and growth directions specification in Arabidopsis petals ^44^. In contrast, the generation of developmental boundaries within floral organs such as petals and a possible involvement of auxin in this process have been scantily investigated.

The flowers of *Hibiscus trionum* constitute an ideal system to study petal patterning because at anthesis mature corollas display a macroscopically visible bullseye characterised by flat, striated and iridescent anthocyanin-rich proximal cells and conical, smooth, flavonol-rich distal cells, separated by a narrow band of smooth tabular boundary cells ^1,2^.

Recently, the emergence of boundary cells has been directly tracked in *Hibiscus trionum*. Before the typical features (pigmentation, cuticle deposition and iridescence, characteristic cell shapes) of the bullseye become detectable, Hibiscus petals are pre-patterned along the proximo-distal axis. A developmental boundary separating the proximal and the distal petal domains, preluding the bullseye patterning of the adaxial epidermal cells, is set earlier than the so-called stage 1 (S1), during the pre-patterning phase (stage 0) that we defined and divided in the stages 0a, 0b, 0c and 1 (S0a, S0b, S0c and S1) ^2,11^. In the distal region, cells are more actively proliferating compared to the proximal region, which show elongated cells and less cell division figures. The cells that later will constitute the bullseye boundary show the largest cell area and are invariably positioned at one third from the petal base in *Hibiscus trionum* during the petal pre-patterning^11^.

Recent computational simulations found that most gene regulatory networks (GRNs) evolved *in silico* to divide the petal surface in two contrasting regions also produce a third cell type at the bullseye boundary^45^. Experimental observations of developing Hibiscus petals suggest that a mobile, non-cell autonomous signal moves from the petal base and triggers the formation of such bullseye boundary cells ^11^. However, how the bullseye boundary is established during the pre-patterning phase is not known and the identity of the patterning signal has not been uncovered. Moreover, the molecular mechanisms that drives epidermal cells to adopt contrasting behaviours—such as distinct growth regimes and differentiation programs—on either side of this boundary during the patterning phase remain to be elucidated.

In Arabidopsis, inter-sepal auxin maxima in the floral meristem specify where petal primordia initiate outgrowth ^32^. Such maxima are established with the contribution of the auxin influx transporter AUXIN1 (AUX1) aided by AUXIN RESISTANCE4 (AXR4), and of the polar auxin efflux carriers PIN1 of the PIN-FORMED (PIN) family ^32^, with *pin1* weak loss of function mutants showing misshaped petals ^46^. Beyond petal initiation, auxin signalling regulates petal growth and morphogenesis, by coordinating cell proliferation with expansion in Arabidopsis ^47^, promoting petal spur elongation in *Aquilegia* ^48^ and controlling size, shape and differential epidermal growth of ray floret ligules in *Gerbera* ^49^. However, whether auxin contributes to the early pre-patterning events that partition the petal surface into distinct developmental territories remains to be investigated.

Here, we use the bullseye of *H. trionum* petals to investigate whether auxin acts as a mobile signal patterning the petal epidermis. Specifically, we test if auxin could (i) induce bullseye boundary formation during the pre-patterning phase and/or (ii) control cell behaviour in the distinct bullseye domains during the later patterning phase. We found that auxin does not specify where and when the bullseye boundary is initiated in Hibiscus petals; rather it regulates cell behaviour in the distal region at later developmental stages, determining the final proportions and appearance of the bullseye.

## RESULTS

### Auxin transcriptional response across the adaxial epidermis does not match bullseye pre-pattern emergence

Auxin gradients are interpreted by plant tissues as coordinate systems to establish growth axes, to specify cell behaviours and cell identities, and set developmental boundaries that precede the generation of patterns ^21,30,50,51^. To test whether an auxin gradient could contribute to bullseye boundary specification during the pre-patterning phase ^11^, we generated a *H. trionum* transgenic line (*DR5v2::ntdTOM*) where the production of the *ntdTOMATO* fluorescent protein is under control of the *DR5v2* synthetic promoter ^52^. After verifying the functionality and responsiveness of this construct in *H. trionum* (**Suppl. Fig. 1a, b**), we imaged the adaxial epidermis of petal primordia during the pre-patterning phase (stage 0a, 0b, 0c and 1) (**Fig. 1a**) when a pigmented bullseye is still undetectable ^11^ and quantified the nuclear *ntdTOMATO* signal as a proxy for auxin transcriptional output (**Fig. 1b**).

**Figure 1.**
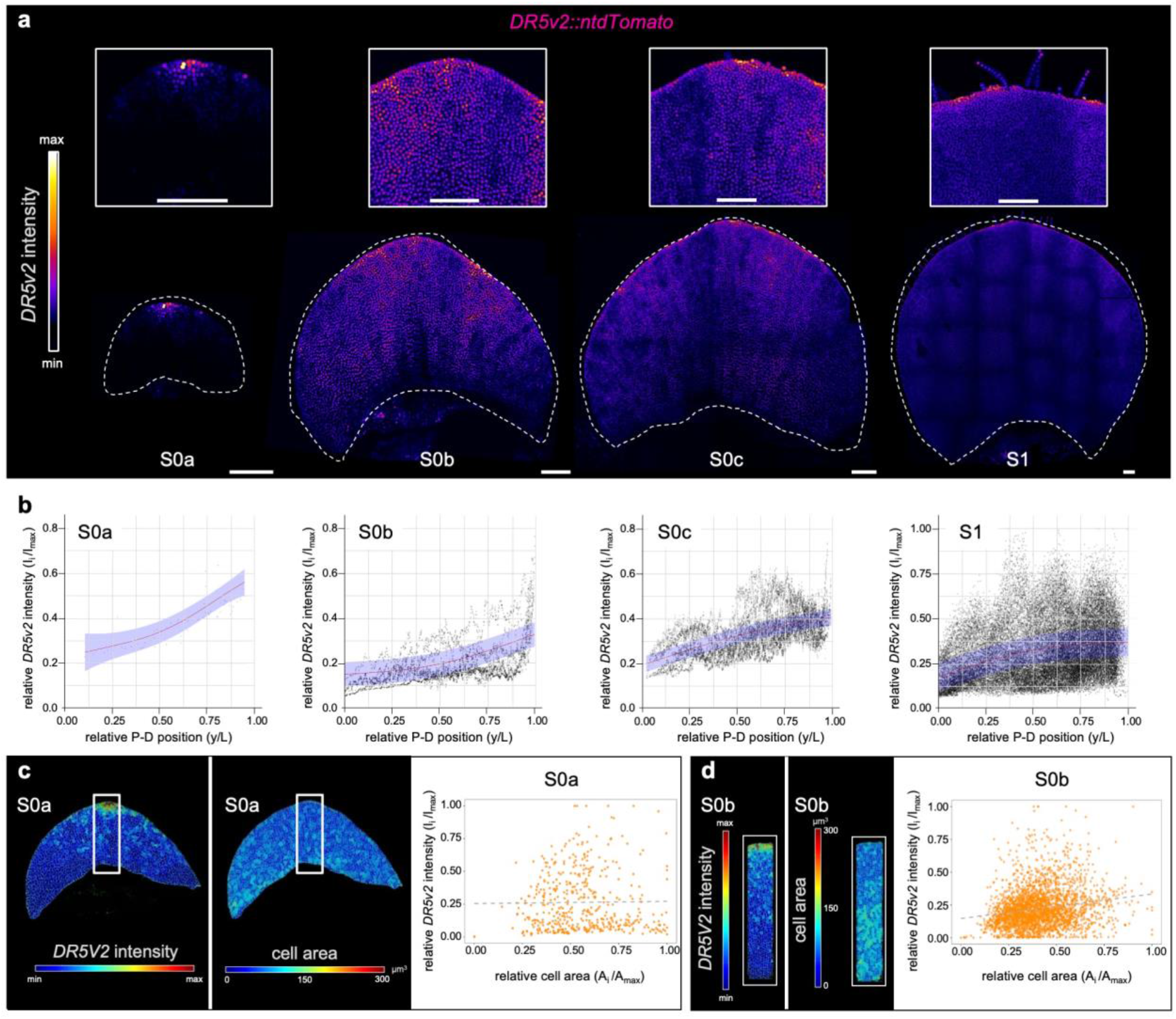
Auxin response is heterogenous across the *H. trionum* petal epidermis but does not predict bullseye pre-pattern emergence. (**a**) Distribution of the auxin transcriptional response in maximum intensity projections of LSCM images across the adaxial epidermis of stage 0 a (S0a) to stage 1 (S1) petal primordia carrying the *DR5v2::ntdTomato* construct. Above are zoom-ins of the most distal regions of each whole petal primordia shown below. *DR5v2::ntdTomato* intensity is false-coloured as indicated in the relative heatmap scale on the left. Scalebars = 100 µm. (**b**) Distribution plots of *DR5v2::ntdTomato* relative intensity (I_i_/I_max_) quantified along the proximo-distal axis y/L at stages S0a-S1. Points represent rolling-average fluorescence intensities calculated within each sample using a window of 10 measurements, red lines show the mean profiles estimated for each stage (generalised additive mixed model GAMM) with smooth effect of relative position and a random intercept for sample. Blue shading indicates approximate pointwise 95% C.I. around the fitted mean profiles (estimate ±1.96 standard errors). For each stage, samples n=6. Biological replicates N=3 (**c**) Spatial distribution maps of *DR5v2::ntdTomato* relative intensity (left panel) and cell areas (middle panel), and their correlation (right panel) at S0a. Pearson’s correlation coefficients: r = 0.367 (approximate unadjusted 95% CI: −0.075–0.688; Holm-adjusted P = 0.0811). (**d**) Spatial distribution maps of *DR5v2::ntdTomato* relative intensity (left panel) and cell areas (middle panel), and their correlation (right panel) at S0b. Pearson’s correlation coefficients: r = 0.246 (approximate unadjusted 95% CI: 0.033–0.437; Holm-adjusted P = 0.0662). In (c) and (d), two-sided one-sample t-tests on sample-specific Fisher-transformed correlations used 5 independent samples per stage.

The strongest *DR5v2* signal was recorded at Stage 0a, as a central focus at the distal tip of the petal epidermis (**Fig. 1a, b**). At stage 0b secondary foci appear distally on the margin of the petal epidermis and in the prospective vein initials. At stage 0c the auxin response relative signal spreads even more along the margin of the petal and across the petal epidermis but remains mostly undetectable in the proximal region. At stage 1, the auxin response remains heterogenous across the developing petal blade, with the highest relative auxin response in the distal region, along the petal edge and with high signal where the incipient vascular cells are. The *DR5v2* signal still gradually decreases toward the proximal petal region (**Fig. 1a, b**). We recorded a similar *DR5v2* expression pattern using a different transgenic *H. trionum* line carrying the *DR5v2* promoter fused to the coding sequence of GFP (*DR5v2::3xGFP*) (**Suppl. Fig. 1c**). These analyses show that *DR5v2* response in Hibiscus petals closely resemble the one observed with *DR5::GUS* and *DR5::GFP* reporter lines in Arabidopsis petals ^44,53^ and indicate that a heterogenous auxin response precedes the emergence of a visible bullseye.

As a band of large anisotropic cells emerges during the pre-patterning phase where the bullseye boundary between purple and cream regions will later appear ^11^, we investigated whether the intensity of the *DR5v2* signal correlates with the cell size distribution that characterises prepatterned petal primordia. We measured the area of petal epidermal cells before prepattern emergence (stage 0a) and during the pre-patterning phase (stage 0b) and tracked the *DR5v2* expression simultaneously (**Fig. 1c, Supp. Fig. 1 d-i**). The analysis shows that at S0a the auxin response maximum localizes at petal tip while cell size across the epidermis is homogenous (**Fig. 1c; Supp. Fig. 1 d-f)**, at S0b the auxin maxima localize on the petal border, and cell heterogeneity appears proximally (**Fig. 1c**; **Fig. 1c, Supp. Fig. 1 g-i)**. In both cases, we found no significant correlation between auxin transcriptional response (*DR5v2* intensity) and pre-pattern (cell area distributions) after correction for multiple testing (mean within-sample correlation: S0a, 0.367, Holm-adjusted P = 0.0811; S0b, 0.246, Holm-adjusted P= 0.0662; n=5 per stage; **Fig. 1c**) suggesting that *DR5v2* intensity does not predict cell area at S0a and S0b. However, from S0c onward, the high level of auxin signalling in the distal region coincides with regions of high cell proliferation. In contrast, the *DR5v2* response is barely detectable in the proximal domain where growth now proceeds via cell expansion instead of division ^11^.

### Interfering with auxin distribution affects petal shape and size but does not prevent bullseye boundary formation

The observed *DR5v2::ntdTOM* expression during the petal pre-patterning stages led us to hypothesize that the emergence of the bullseye boundary is not a direct read-out of the auxin response. To test this hypothesis, we used chemical treatments that perturb auxin distribution in developing petal primordia. We applied different concentrations of the polar auxin transport inhibitor 2,3,5-triiodobenzoic acid (TIBA) ^54,55^ or the synthetic auxin analogue 2,4-Dichlorophenoxyacetic acid (2,4-D) ^56^ to young buds bearing unpatterned petal primordia (< stage 1) and allowed them to reach maturity. We then recorded the phenotype of open flowers (stage 5) (**Fig. 2**). We found that TIBA treatments never prevented bullseye formation (45/45 obs. in total, n ≥ 10 for each treatment concentration) (**Fig. 2a**). However, treated flowers produced smaller and narrower petals in a concentration-dependent manner at higher dosage treatments (**Fig. 2d, e; Suppl. Fig. 2a**), with both proximal and distal regions affected similarly (**Fig. 2f, g; Suppl. Fig. 2a**) thereby resulting in unaltered bullseye proportions (**Fig. 2h**, **Suppl. Fig. 2a**). Similarly, the ectopic applications of 2,4D, a synthetic auxin which mimics its activity but is scarcely transported by PINs and instead accumulates within the cells, did not prevent bullseye formation (75/75 obs., n≥10 for each treatment concentration) (**Fig. 2b**). Like TIBA treatments, 2,4D applications negatively impacted petal growth in both length and width in a dose dependent manner compared to control flowers (mock treated) (**Fig. 2i, j**). The distal petal region was more affected than the proximal domain (**Fig. 2k, l; Suppl. Fig. 2b**): at lower 2,4-D concentrations, the length of the distal region was more impacted than the proximal domain, showing that the distal region is more responsive to the treatment (**Suppl. Fig. 2b**). The absolute position of the bullseye boundary along the petal proximo-distal axis was unaffected for low to mid (2.5, 5, 10 mM) 2,4D concentration treatments, as the petal proximal domain was only shorten when higher (50 and 100 mM) 2,4-D concentrations were applied (**Fig. 2k, l; Suppl. Fig. 2b**). Therefore, bullseye proportions were not conserved but increased as flower dimensions decreased (**Fig. 2m**). Notably, at high (50 and 100 mM) 2,4-D concentration, most epidermis cells in the distal domain failed to acquire their characteristic conical shape and remained flat (conical cells were only detected near the petal distal margin, **Fig. 2c**), with the distal petal region retaining the green, immature appearance of earlier developmental stages (**Fig. 2b, c**). Hence, both growth and cell differentiation are severely affected in the distal bullseye region while the proximal petal domain seems less sensitive to chemical alterations of auxin cellular levels.

**Figure 2.**
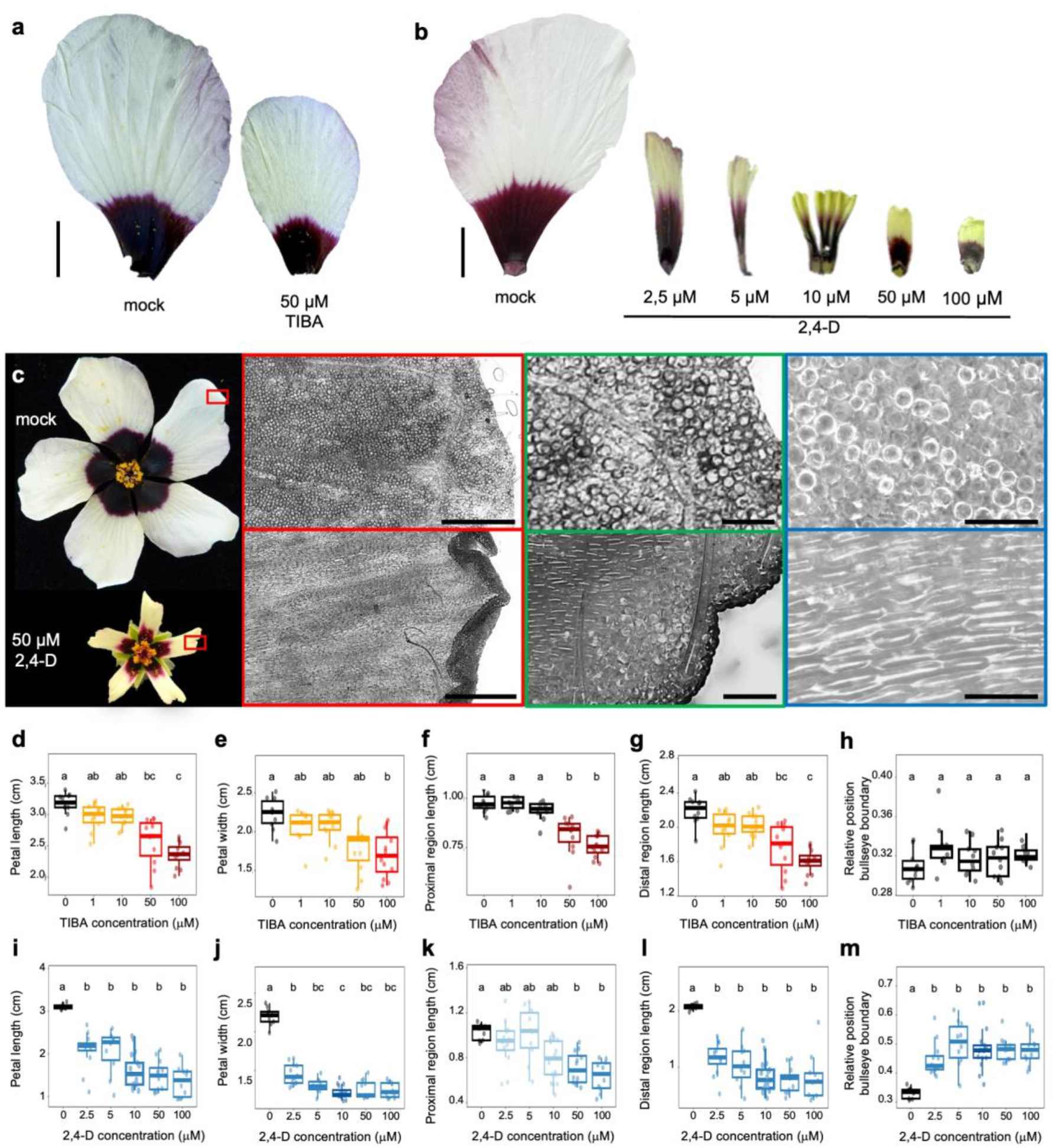
Application of an auxin transport inhibitor or a low-mobility auxin analogue modifies petal size and bullseye proportions. *H. trionum* petals of mature flowers (S5) after treatment with (**a**) mock and 50 µM TIBA or (**b**) mock and 2.5, 5, 10, 50 and 100 µM 2,4-D. Scalebars = 1cm. (**c**) *H. trionum* mature flowers (S5) treated with mock and 50 µM 2,4-D respectively, with red square indicating the distal petal area of the zoom-ins shown in panels alongside. 2,4D treatment negatively interferes with normal conical cell development: conical cells are only found near the petal top margin when plants are treated with 50 µM 2,4-D (green panel) and conical cells are largely replaced with flat tabular cells throughout most of the distal region (blue panels). Scale bars in red panel = 500 µm, in green panel= 100 µm, in blue panel = 100 µm. (**d**) Petal length, (**e**) petal width, (**f**) length of the proximal region, (**g**) length of the distal region and (**h**) relative position of the bullseye boundary in *H. trionum* petals of mature flowers (S5) following TIBA applications at the indicated treatment concentrations and in mock control. (**i**) Petal length, (**j**) petal width, (**k**) length of the proximal region, (**l**) length of the distal region and (**m**) relative position of the bullseye boundary in *H. trionum* petals of mature flowers (S5) following 2,4-D applications at the indicated treatment concentrations and in mock control. For all boxplots, each point represents the mean of ≥3 petals from one flower. For TIBA, n = 10, 10, 11, 11 and 12 flowers for mock, 1, 10, 50 and 100 µM, respectively. For 2,4-D, n = 9, 12, 10, 17, 12 and 10 flowers for mock, 2.5, 5, 10, 50 and 100 µM, respectively. Values reported are mean ± SD; percentage changes are relative to the corresponding mock mean. For all tests, flower-level measurements were analysed using Kruskal–Wallis tests, with Holm correction within each treatment. Two-sided pairwise Wilcoxon rank-sum tests were performed with Bonferroni correction across all pairwise comparisons within each treatment. Groups sharing at least one letter were not significantly different from each other; groups sharing no letters differed significantly, with adjusted p-value <0.05.

Taken together, our results reveal that bullseye boundary formation is insensitive to treatments impairing auxin distribution but that perturbations of the endogenous auxin content influence cell behaviour (growth and differentiation) differentially across the petal epidermis, with the two bullseye domains displaying distinct sensitivities to perturbations of auxin cellular contents.

### Genetic modifications of auxin distribution do not prevent boundary formation but impact cell behaviour differentially across the petal surface

In Arabidopsis petals, the transcription factors JAGGED (JAG) and PETAL LOSS (PTL) contribute to petal morphogenesis from early developmental stages by interacting with the auxin response and impacting the *DR5* map across the petal ^32,44^. We identified six *PTL* homologs (*HtPTL1*-6) and three *JAG* homologs (*HtJAG1, HtJAG2, HtJAG3*) in the *H. trionum* genome ^57^. Based on the transcriptomic data from unpatterned S1 *Hibiscus* petal primordia ^58^, all paralogs expressed in the S1 petals share an expression domain similar to their Arabidopsis counterparts: the *HtPTL* genes are preferentially transcribed in the proximal region (**Suppl. Fig. 3a**) while the *HtJAG* genes are more expressed in the distal petal domain (**Suppl. Fig. 3b**).

Therefore, we overexpressed in *H. trionum HtPTL1* and *HtJAG1* to assess the impact of modifying the spatio-temporal auxin signalling response on petal development and bullseye pattern formation (**Fig.3; Suppl. Fig. 3d, i**). We found that the constitutive overexpression of *HtPTL1* (*HtPTL1* OE) reduces the area of stage 5 petals (**Fig. 3a, b; Suppl. Fig. 3c, e-f**) by significantly diminishing the distal area regions (mean distal region area of *HtPTL1* OE is 26% smaller than WT; 3.2 ± 0.5 cm² for *HtPTL1* OE vs. 4.3 ± 0.7 cm² for WT; t = 4.344, df = 16.16, adjusted p-value = 0.00148) without abolishing bullseye formation (obs: 100% of flowers from 4 transgenic lines, n=40 flowers/transgenic line). Misexpressing *HtPTL1,* thus, triggers an increase in bullseye proportions (**Fig. 3c**) due to growth reduction of the distal domain (**Suppl. Fig. 3c**). The smaller size of the distal region is likely due to a reduction in cell number, rather than a reduction in cell size as distal and proximal cells in *HtPTL1* OE are similar to wild-type (**Fig. 3d, e).** Such reduction in cell proliferation is consistent with the known role of PTL as a growth repressor in *Arabidopsis* 59,60_._

**Figure 3.**
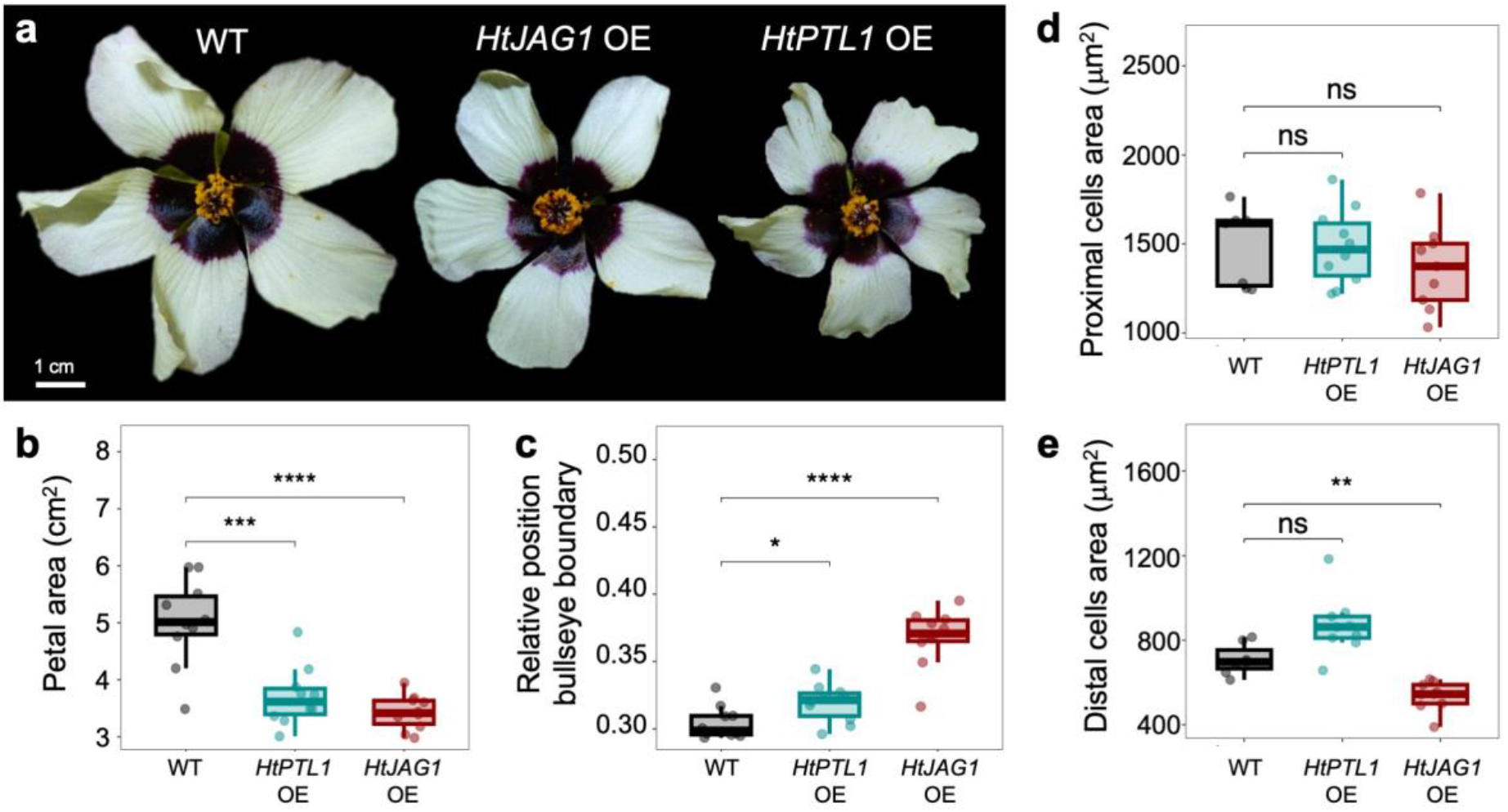
Constitutive overexpression of *HtJAG1* and *HtPTL1* alters distal petal growth without affecting bullseye boundary formation. (**a**) Representative *Hibiscus trionum* flowers (S5) of wild type (WT), *HtJAG1* OE and *HtPTL1* OE lines. Scalebar = 1cm. (**b**) Petal area, (**c**) relative position of the bullseye boundary and cell area in the (**d**) proximal and (**e**) distal petal region of *H. trionum* mature flowers (S5) of wild type (WT), *HtJAG1* OE and *HtPTL1* OE lines. Total petal area is reduced by 26.2% in *HtPTL1* OE compared to WT (3.70 ± 0.52 versus 5.02 ± 0.76 cm²; Student’s two-sample t-test, t = 4.497, df = 18, P = 0.000279) and by 31.7% in *HtJAG* OE (3.43 ± 0.30 versus 5.02 ± 0.76 cm²; Welch’s t-test, t = 6.114, df = 11.76, P = 5.69 × 10⁻⁵); the bullseye boundary was shifted upwards along the proximo-distal axis of the petal by 4.9% in *HtPTL1 OE* (0.32 ± 0.01 versus 0.30 ± 0.01; Student’s two-sample t-test, t = −2.495, df = 18, P = 0.0226) and by 20.8% in *HtJAG1* OE petals compared to WT (0.37 ± 0.02 versus 0.30 ± 0.01; Student’s two-sample t-test, t = −8.010, df = 18, P = 2.41 × 10⁻⁷); n=10 flowers per genotype. ns = not significant (adjusted p-value ≥ 0.05), * = adjusted p-value < 0.05, ** = adjusted p-value < 0.01, *** = adjusted p-value < 0.001.**** = adjusted p-value < 0.0001. Petal epidermis cells did not significantly differ in size when *HtPTL1* was constitutively overexpressed (Proximal cells: 1487 ± 212 µm^2^ in *HtPTL1 OE* vs. 1488 ± 221 µm² in WT; two-sided Welch’s t-test, t = 0.042, df = 12.71, Holm-adjusted P = 0.9674; n = 10 and 7, respectively; Distal cells: 871 ± 142 µm^2^ in *HtPTL1 OE* vs. 709 ± 75 µm² in WT; two-sided Welch’s t-test, t = −2.935, df = 12.56, Holm-adjusted P = 0.0599; n = 9 and 7, respectively). Proximal cells did not significantly differ in size when *HtJAG1* was constitutively overexpressed but distal cells were significantly smaller compare to WT but distal cells were 24% smaller in *HtJAG1* OE compared to WT (Proximal cells: 1366 ± 235 µm^2^ in *HtJAG1 OE* vs. 1488 ± 221 µm² in WT; t = 1.070, df = 13.40, Holm-adjusted P = 0.6112; n = 9 and 7, respectively; Distal cells: 537 ± 70 µm^2^ in *HtJAG1 OE* vs. 709 ± 75 µm²; t= 4.686, df = 12.61, Holm-adjusted P = 0.00323; n = 9 and 7, respectively). For all analyses, values are mean ± SD; percentage changes are relative to the corresponding wild type mean. For boxplots in b-c, each point represents the mean of ≥3 petal from one flower, for boxplots in d-e each point represents the mean of ≥10 cells from one petal. Student’s or Two-sided Welch’s t-tests compared each OE line with WT, with Holm correction across all comparisons.

On the other hand, the constitutive overexpression of *HtJAG1* (**Suppl Fig. 3i**) yielded plants with elongated bracts (**Suppl Fig. 3g-h**). *JAG* overexpression promotes cryptic bract outgrowth in Arabidopsis ^61,62^, supporting the biological activity of the *HtJAG1* OE construct. We found that constitutive overexpression of *HtJAG1* does not prevent the formation of the bullseye, which was recorded in the totality of the analysed flowers (100/100 flowers from 5 transgenic lines, n=20 flowers/transgenic line) at stage 5 (**Fig. 3a**). However, *HtJAG1* OE plants produce flowers where the overall area of the petal is shrunk (**Fig. 3b**). We quantified the dimensions of the bullseye region and found that *HtJAG1* OE smaller petals are almost exclusively due to a reduction of the distal region (mean distal region area of *HtJAG1* OE is 35% smaller than WT; 2.8 ± 0.3 cm² for *HtJAG1* OE vs. 4.3 ± 0.7 cm² for WT; t = 6.612, df = 11.62, adjusted p-value = 0.000117) while the size of the proximal region is maintained (0.6 ± 0.1 cm² for *HtJAG1* OE vs. 0.7 ± 0.1 cm² for WT; t = 2.297, df = 13.21, adjusted P = 0.0772) (**Suppl Fig. 3e-f**), hence changing the proportion of the bullseye (**Fig. 3c**) in a way that is reminiscent of the phenotypes observed upon 2,4-D lower concentration treatments (**Fig. 2**). The smaller distal region of *HtJAG1* OE flowers is due, at least in part, to a reduction in conical cell size while the dimensions of flat tabular cells remain unchanged in the proximal domain (**Fig. 3d-e**).

Overall, altering the auxin response map through genetic manipulation of *HtPTL1* and *HtJAG1* expression further corroborates the observations that auxin signalling differentially modulates petal growth without interfering with boundary formation. Our results show that, while *HtJAG1* and *HtPTL1* have distinct expression domains, overexpression of either factor preferentially affects cell behaviour in the distal region of the petal, where the strongest auxin-transcriptional response is recorded. This suggests that the correct development of the distal region is particularly sensitive to the crosstalk between JAG and PTL that helps modulate auxin distribution.

### Genetic perturbation of auxin homeostasis affects petal initiation, number and growth, but does not interfere with bullseye formation

To understand how auxin signalling impacts cell behaviour in the distal domain of the bullseye, we examined transcriptomic data from S1 and S2 petal primordia ^45,58^ and extracted auxin-related genes preferentially expressed in the distalmost petal region. This singled out auxin homeostasis genes such as group II *GRETCHEN HAGEN 3* genes, that code for acyl acid amido synthetases that catalyse the Asp and Glu aminoacid conjugation to IAA and IAA irreversible inactivation as well as *IAA CARBOXYL METHYLTRANSFERASE1* (*IAMT1*), which encodes an enzyme responsible for the methylation of indole-3-acetic acid (IAA) and the production of inactive methyl-IAA (Me-IAA) (**Suppl. Fig. 4 a**).

To further investigate the role of auxin in petal patterning by reducing the IAA content while possibly overcoming gene redundancy, we generated constitutive overexpression lines for *HtIAMT1* as well as three of the most highly differentially expressed (|log₂FC| > 2, p.adj<0.05) group II *GH3.1* homologs (*HtGH3.1a, b* and *c*). After validating the transgenic overexpression lines (**Suppl. Fig. 4f-i**), we analysed the phenotypic effects of these genetic manipulations. A clear bullseye was present in all analysed flowers (100% of analysed samples, n>10 for each transgenic line) (**Fig.4 a; Suppl. Fig. 4j)** further supporting the idea that bullseye pattern establishment is robust to genetic modification of auxin homeostasis and levels. However, most lines show obvious changes in petal size alongside a plethora of altered flower phenotypes (flower sterility, “double-flowers”, altered specification of petal conical cells, altered specification of abaxial-adaxial petal polarity) (**Fig. 4a**) that resemble those reported when homologous and related genes are altered in Arabidopsis and other species flowers ^63^.

**Figure 4.**
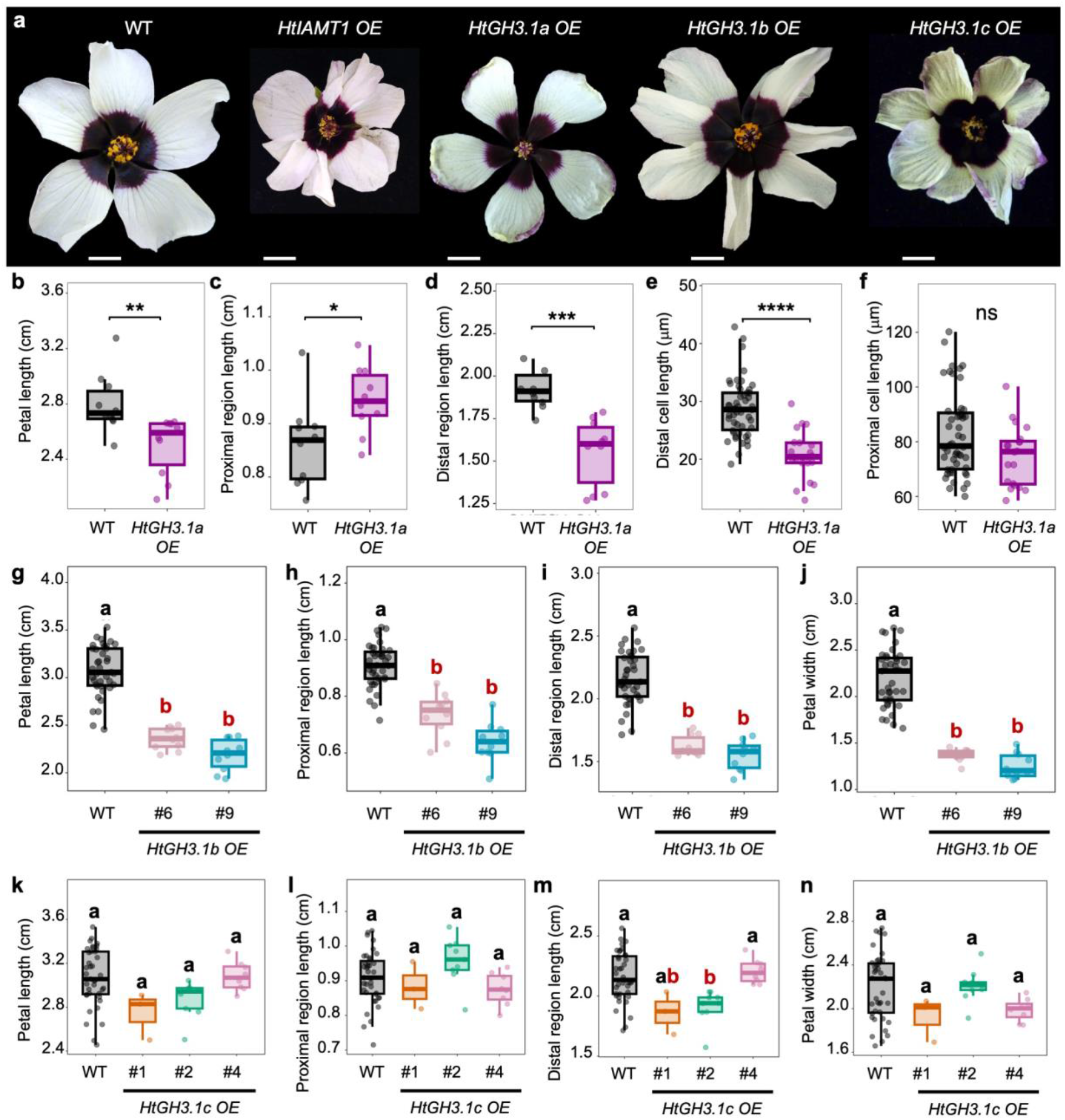
Genetic disruption of auxin homeostasis uncouples petal initiation and growth from bullseye formation. (**a**) Representative *Hibiscus trionum* flowers (S5) of wild type (WT), *HtIAMT1* OE*, HtGH3.1a* OE*, HtGH3.1b* OE, and *HtGH3.1c* OE lines. Scalebars = 1 cm. (**b**) Petal length, (**c**) length of the proximal petal region, (**d**) length of the distal petal region, length of (**e**) distal and (**f)** proximal cells in *H. trionum* mature flowers (S5) of wild type (WT) and *HtGH3.1a* OE line. Compared to WT, *HtGH3.1a* OE line #1 showed a 5% reduction in median petal length (2.592 [IQR: 0.298] versus 2.734 [IQR: 0.200] cm; two-sided Wilcoxon rank-sum test with continuity correction, W=93, P=0.001315); a 10% increase in mean proximal region length (0.944 ± 0.062 versus 0.860 ± 0.080 cm; mean ± SD; two-sided Student’s t-test, t{18}=-2.610, P=0.01772) and a 20% reduction in mean distal region length (1.555 ± 0.197 versus 1.940 ± 0.147 cm; mean ± SD; two-sided Student’s t-test, t{18}=4.951, P=1.032 10^-4^). Distal conical cells were 28% shorter in *HtGH3.1a* OE line #1 (20.8 ± 4.2 µm in *HtGH3.1a* OE vs. 28.7 ± 4.9 µm in WT; mean ± SD; two-sided Student’s t-test, t{68}=-6.415, P=1.579×10^-8^) while the length of the proximal flat striated cells did not significantly differ between *HtGH3.1a* OE line #1 and WT (median [IQR]: 76.4 [15.7] µm in *HtGH3.1a* OE vs. 78.4 [20.6] µm in WT; two-sided Wilcoxon rank-sum test with continuity correction, W=354, P=0.05854). For panels b–d, each point represents one flower-level value, with (n=10) observations per genotype. For panels e-f, each point represents the mean of multiple cell-length measurements within one petal. All reported (P) values are unadjusted. ns = not significant (p-value ≥ 0.05), * = p-value < 0.05, ** = p-value < 0.01, *** = p-value < 0.001.**** = p-value < 0.0001. (**g**) Petal length, (**h**) length of the proximal petal region, (**i**) length of the distal petal region, (**j**) petal width in *H. trionum* mature flowers (S5) of wild type (WT) and *HtGH3.1b* lines. Compared to WT, mean petal length was reduced by 23% and 29% in *HtGH3.1b* OE line #6 and line #9, respectively (Holm-adjusted P=7.09×10^-7^ and P=3.29 x10^-8^, respectively) while petal width was reduced by 38% in *HtGH3.1b* OE line #6 and 43% in line #9 (Holm-adjusted P=6.55 x10^-8^ for both comparisons). Both proximal and distal regions were shortened: the mean proximal region length was reduced by 19% in *HtGH3.1b* OE line #6 and 30% in line #9 (Holm-adjusted P=1.16 x10^-4^ and P=2.59 x10^-7^, respectively). The mean length of the distal region was reduced by 25% in *HtGH3.1b* OE petals line *#6* and 28% in line #9 (Holm-adjusted, P=7.71 x10^-7^ and (P=6.55 x10^-8^, respectively). For panels g-j, each point represents one flower-level mean calculated from measurements of 3-5 petals, with n=39 for WT and n *≥* 10 for each transgenic line. Overall differences were assessed using permutation Kruskal–Wallis tests, with Holm correction. Pairwise comparisons used two-sided exact Wilcoxon–Mann–Whitney tests, with Holm correction across comparisons. Groups sharing at least one letter were not significantly different; groups sharing no letters differed at adjusted (P<0.05). (**k**) Petal length, (**l**) length of the proximal petal region, (**m**) length of the distal petal region, (**n**) petal width in *H. trionum* mature flowers (S5) of wild type (WT) and *HtGH3.1c* OE lines. Differences in petal length and width between genotypes were not statistically significant (length: Holm-adjusted P=0.652, P=0.608, and P=1.00, for line *HtGH3.1c* OE #1, 2 and 4, respectively; width: Holm-adjusted P=1.00, for all lines *HtGH3.1c* OE #1, 2 and 4. The length of the proximal region did not significantly differ between genotypes (Holm-adjusted P=1.00 for all three comparisons) but the length of the distal region was significantly reduced in *HtGH3.1c* OE line #2 (Holm-adjusted P=0.0183). For panels k-n, each point represents one flower-level mean calculated from measurements of 3-5 petals; with n=39 for WT and n=3 for line #1, n=9 for line #2, and n=8 for line #4. Overall differences were assessed using permutation Kruskal–Wallis tests, with Holm correction. Pairwise comparisons used two-sided exact Wilcoxon–Mann–Whitney tests, with Holm correction across comparisons. Groups sharing at least one letter were not significantly different; groups sharing no letters differed at adjusted (P<0.05).

Specifically, we found that overexpressing *HtIAMT1,* causes “double flowers” and flower fasciation (**Fig. 4a; Suppl. Fig. 4j**) but does not alter the geometry of the bullseye (**Suppl**. **Fig. 4n**) because the size of proximal and distal regions are equally scaled down, a phenotype reminiscent of the effects of the TIBA petal treatments (**Fig. 2a)**. When we overexpressed *HtGH3.1a*, we retrieved two independent transgenic lines. One could not reach anthesis, with flowers aborting within the buds before reaching S5 (**Suppl. Fig. 4b**). For the line with flowers reaching anthesis (**Fig. 4a; Suppl. Fig. 4j**), we recorded shorter petals (**Fig. 4b; Suppl. Fig. 4c-d**) caused by a 20% reduction of the petal distal region (1.56 ± 0.20 cm in *HtGH3.1*a OE versus 1.94 ± 0.15 cm in WT; mean ± SD; two-sided Student’s t-test, (t{18}=4.951, p-value = 0.0001032), while the length of the proximal region increased slightly (0.94 ± 0.06 in *HtGH3.1*a versus 0.86 ± 0.08 cm in WT; mean ± SD; two-sided Student’s t-test, t{18}=-2.610, p-value = 0.01772) (**Fig. 4c, d**). The shrinkage of the distal region, and increase in bullseye size, is due, at least in part, to a reduction in conical cell size while proximal tabular cells remain unchanged (**Fig. 4e, f**). These distal petal cells also fail to fully acquire their characteristic conical shape, resembling the phenotype observed when WT plants were treated with 2,4-D and this likely causes the distal region of the petal to appear thicker than WT and upward curled (**Suppl. Fig. 4c-e)**. Three lines ubiquitously overexpressing *HtGH3.1b* consistently showed a flower phenotype: we observed petal fasciation or polypetalous flowers (**Fig. 4a**) and shorter, narrower petals compared to WT due to a reduction of both the proximal and distal regions (**Fig. 4g-j**), with no significant alteration in bullseye proportions (**Suppl. Fig. 4l**). We also recorded polypetalous flowers when *HtGH3.1c* was overexpressed (**Fig. 4a; Suppl. Fig. 4j**) but petal length and width were not significantly affected (**Fig. 4k,n**). Bullseye proportions were significantly increased in one line only (line #2), due to a shortened distal region (**Fig. 4 l,m; Suppl. Fig. 4m**), again suggesting that auxin depletion through missexpression of *HtGH3.1c* can affect growth of the distal domain.

Altogether, we found that endogenously altering auxin homeostasis through overexpression of group II *GH3.1* and *IAMT1* homologs does not prevent bullseye formation but disrupt the mechanisms controlling petal number, size and shape as well as distal cell differentiation. Notably, when we overexpressed group II *GH3.1* or *IAMT1* genes, normally preferentially expressed in the distal region at S1-S2 (**Suppl. Fig. 4a**), we recorded the strongest phenotypic defects (impaired growth and cell differentiation) in the distal region. Hence, tweaking auxin homeostasis *in planta* corroborates our previous observations that the distal region of the petal relies on fine balances of auxin levels to grow and differentiate correctly.

### *HtTCP4* genes control bullseye proportions by promoting auxin biosynthesis in the distal region of the petal

*HtTCP4.1* and *HtTCP4.2*, two homologs of the TCP4 transcription factor are preferentially expressed in the distal region of S1 petal primordia ^11^. We previously reported that plants overexpressing *HtTCP4.1* produce flowers with enlarged bullseye due to increased cell proliferation at the base of the stage 0a petal, without altering the bullseye boundary pre-patterning nor the mature petal overall area ^11^. As the high auxin transcriptional response in the distal region overlaps with *HtTCP4* genes expression, we wondered whether HtTCP4.1 and HtTCP4.2 act by boosting auxin signalling to promote cell proliferation in the distal region of the petal. To investigate this, we introduced the *DR5v2::ntdTOM* construct into the *HtTCP4.1* OE line ^11^ and imaged stage 0a petals. We observed that a modified relative auxin transcriptional response profile, with a higher *DR5v2* fluorescence recorded in the *HtTCP4.1* OE distal region compared to wild-type plants (**Fig. 5a, b**), consistently with a role for HtTCP4 homologs in promoting auxin signalling during Hibiscus petal development.

**Figure 5.**
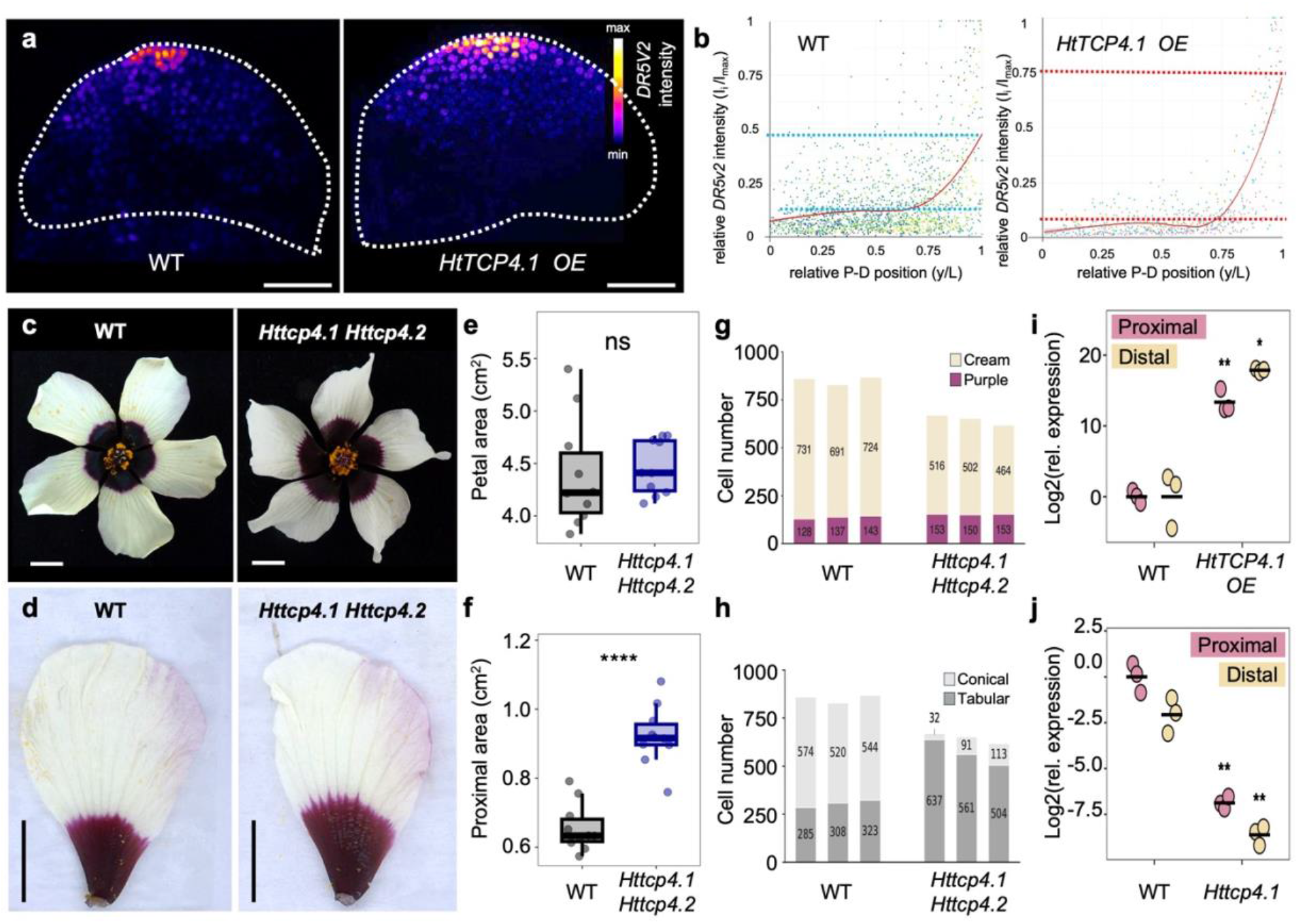
TCP4 homologs stimulate *HtYUCCA3* expression to promote auxin biosynthesis and cell proliferation in the distal petal region of *H. trionum* flowers. (**a**) Spatial distribution map of the auxin transcriptional response visualized by LSCM across the adaxial epidermis of stage 0a (S0a) petal primordia expressing *DR5v2::ntdTomato* in WT and *HtTCP4.1 OE*. *DR5v2::ntdTomato* intensity is false-coloured as indicated in the relative heatmap scale on the right. Scalebars = 50 µm. (**b**) Distribution plots of *DR5v2::ntdTomato* relative intensity (I_i_/I_max_) quantified along the proximo-distal axis y/L at stages S0a of *DR5v2::ntdTomato* in *Wt* and in *HtTCP4.1 OE*. Points represent individual measurements, red lines show the mean profiles estimated for each stage (generalised additive mixed model) with smooth effect of relative position and a random intercept for sample. Gray shading indicates approximate pointwise 95% C.I. around the fitted mean profiles (estimate ±1.96 standard errors). For each stage samples n=5; biological replicates N=3. Dashed lines (light blue for *Wt* and red for *HtTCP4.1 OE*) indicate the fitted relative intensities at y/L = 0.50 and 1.00. (**c**) Representative *Hibiscus trionum* flowers (S5) of wild type (WT) and *Httcp4.1 Httcp4.2* double mutant. Scalebars = 1cm. (**d**) Representative *Hibiscus trionum* petals (S5) of wild type (WT) and *Httcp4.1 Httcp4.2* double mutant. Scalebars = 1cm. (**e**) Petal areas of WT and *Httcp4.1 Httcp4.2* double mutant. Petal area was similar in both genotypes (4.5 ± 0.3 cm^2^ in *Httcp4.1 Httcp4.2* double mutant vs. 4.4 ± 0.5 cm² in WT; adjusted P = 0.800). (**f**) Area of the proximal petal region of WT and *Httcp4.1 Httcp4.2* double mutant. The mean proximal region area was increased by 40.6% in the double mutant compared to WT (0.92 ± 0.09 cm^2^ in *Httcp4.1 Httcp4.2* double mutant vs. 0.66 ± 0.07 cm² in WT; adjusted p-value = 8.31 × 10⁻⁶). For e-f values are means ± SD, percentage changes are relative to the wild-type means. Each point represents one flower mean calculated from measurements of its petals, with n≥10 flowers per genotype. Boxes indicate the interquartile range, horizontal lines mark medians, and whiskers extend to the most extreme observations within 1.5 times the interquartile range. WT is shown in black, *Httcp4.1 Httcp4.2* in blue. Cell counts classified by pigmentation (**g**) or shape (**h**) in individual petals of WT and *Httcp4.1 Httcp4.2* double mutant. Each stacked bar represents cell count for one petal along the proximo-distal axis (numbered 1–3 for each genotype), with purple cells shown in purple and off-white cells in cream. Tabular cells are displayed in dark grey and conical cells in light grey. Segment labels indicate cell counts, and total bar height indicates total cell number along the proximo-distal petal axis. n = 3 petals per genotype. (**i**) qRT-PCR quantification of *HtYUC3* relative transcript abundance (log_2_ relative expression) in proximal and distal tissue of *HtTCP4.1* OE and WT S3 petals. For both petal regions, expression was significantly higher in the overexpression transgenic line compared to WT (proximal, adjusted p-value = 0.00233; distal, adjusted P = 0.0152). (**j**) qRT-PCR quantification of *HtYUC3* relative transcript abundance (log_2_ relative expression) in proximal and distal tissue of *Httcp4.1* mutant and WT S3 petals. For both petal regions, expression was lower in the knockout line (adjusted p-value = 0.00233 for each comparison). For i-j expression was analysed on the log₂ scale using two-sided Welch’s t-tests comparing each transgenic or mutant line with its corresponding WT control within each petal region. P-values were adjusted using the Benjamini–Hochberg procedure across the four comparisons. N=3. * = adjusted p-value < 0.05, ** = adjusted p-value < 0.01.

To test if *HtTCP4* genes promote cell proliferation in the distal bullseye domain, we used CRISPR/Cas9 genome editing to generate double mutants for these genes (**Suppl. Fig. 5f**). We predicted that *Httcp4.1 Httcp4.2* plants might produce smaller flowers with a wild-type proximal petal region but a reduced distal domain due to a lack of cell proliferation in the top petal region. Surprisingly, double mutant individuals yield flowers no different in size to wild-type ones (WT mean petal area = 4.39 ± 0.52 cm^2^, *Httcp4.1-1 Httcp4.2-6* double mutant mean petal area = 4.46 ± 0.26 cm^2^, unpaired two-sample t-test, p-value = 0.723) (**Fig. 5c-e, Suppl. Fig. 5a,b**) but with an enlarged proximal region (proximal region 39% larger in *Httcp4.1-1 Httcp4.2-6* double mutant; mean area of proximal region in WT = 0.66 ± 0 .07 cm^2^; mean area of proximal region in double mutant = 0.92 ± 0.09 cm^2^, Student’s Two-sample t-test p-value <0.0001, **Fig. 5f**) and thus, a bigger bullseye (bullseye size = 21% ± 2% in *Httcp4.1-1 Httcp4.2-6* double mutant vs. 15% ± 0.5% in WT, Welch’s Unpaired t-test, p-value <0.0001, **Suppl. Fig. 5d**). However, careful examinations at the cellular level revealed that the number of distal cells along the proximo-distal axis was significantly reduced in *httcp4.1-1 httcp4.2-6* flowers (31% reduction in cell number with 494 ± 27 cells in double mutant vs. 715 ± 22 cells in WT, Student’s two-samples t-test, p-value<0.0001), while the number of proximal, purple-pigmented cells was unchanged (136 ± 8 cells in WT vs. 149 ± 5 cells in double mutant, p-value = 0.074] (**Fig. 5g, Suppl. Fig. 5e**). This supports the idea that an absence of HtTCP4 activity strongly decreases cell division but only significantly in the distal region, where both paralogs are preferentially expressed. Interestingly, we found that petal length was maintained in the double mutant (petal length = 3.08 ± 0.15 cm in WT vs. 3.15 ± 0.11 cm in *Httcp4.1-1 Httcp4.2-6* double mutant, unpaired two-sample t-test, p-value = 0.263, **Suppl. Fig. 5a**) because purple proximal cells in *Httcp4.1-1 Httcp4.2-6* were significantly longer than equivalent cells in wild type (30% increase in proximal cell length, 65 ± 10 µm in WT vs. 84 ± 15 µm in double mutant, p-value < 0.0001, **Suppl. Fig. 5c**), accounting for the production of a bigger bullseye, and because a large proportion of distal cells were tabular (82-95% of petal cells are tabular in *Httcp4.1-1 Httcp4.2-6* vs. 33-37% in wild type) instead of conical (567 ± 67 tabular cells along the PD axis in the double mutant vs. 305 ± 19 tabular cells along the PD axis in WT, p-value = 0.0028, **Fig. 5h**; **Suppl. Fig. 5e**).

Hence, the absence of HtTCP4 activity impairs both cell division and correct differentiation in the distal petal region, a phenotype reminiscent of those produced by perturbing auxin signalling and homeostasis (**Fig. 2-4**). Whether the increase in cell elongation is a direct effect of HtTCP4.1 and HtTCP4.2 absence or whether this is indicative of independent compensatory mechanisms aiming at conserving petal size remains to be established. Regardless, it is interesting to note how both overexpressing and knocking-out *HtTCP4* genes produce flowers with larger bullseyes but through different mechanisms: an increase in the number of proximal cells in *HtTCP4.1* OE ^11^ or a decrease in distal cell number accompanied with an increase in cell length across the petal surface in the *Httcp4.1 Httcp4.2* double mutant.

In Arabidopsis, TCP4 binds the promoter of *YUCCA5* (*YUC5*), and induces its expression to elicit differential growth powering hypocotyl elongation ^64^. *YUC5* codes for a flavin-containing monooxygenase that catalyses the rate-limiting conversion of the precursor indole-3-pyruvic acid (IPA) into the active auxin Indole-3-acetic acid (IAA) ^65^. To test whether *HtTCP4* genes promote cell proliferation in the distal region by stimulating auxin synthesis there, we searched for differentially expressed *HtYUCCA* genes in the transcriptome of S1 petals ^58^. We found that a homolog of *YUCCA3*, *HtYUCCA3*, is preferentially expressed in the distal petal region where the higher *DR5v2* auxin response is recorded (**Fig.1 a-b; Suppl. Fig. 1c**). We investigated the expression of *HtYUC3* in both *HtTCP4.1* OE and in the *Httcp4.1* mutant and found that it is respectively more and less expressed compared to wild type (**Fig.5 i,j**). These results indicate that *HtTCP4* genes promote the expression of *HtYUC3* and suggest a plausible mechanism through which HtTCP4 regulate cell proliferation by boosting *HtYUC3* transcription in the distal region of the Hibiscus petal, locally promoting auxin anabolism.

Altogether, our results support a model in which auxin contributes to petal initiation and regulation of organ number but is not required to establish bullseye boundary during the pre-patterning phase. Instead, auxin signalling differentially regulates cell behaviour, and thus growth, across the petal surface: high auxin activity in the distal domain accounts for the maintenance of cell proliferation up to stage 4 and correct cell-shape elaboration during the patterning phase, whereas the proximal domain, characterised by a lower auxin response, undergoes an early arrest of cell division by stage 2 and instead grows via expansion of flat tabular cells ^11^.

## DISCUSSION

Extensive research across a range of model species has provided us with a detailed understanding of how flowering and floral organ specification are regulated ^66–68^ but the subsequent processes that pattern floral tissues to generate functional organs and surfaces remain less characterised. Here, we used *Hibiscus trionum* to explore the hypothesis that auxin, a primary phytohormone governing nearly every facet of plant development, functions as a mobile signal initiating bullseye pattern formation on the petal surface. We previously reported that the bullseye boundary is established very early during a pre-patterning phase in *H. trionum* and that petal primordia subsequently experience a 100-fold growth ^11^. Strikingly the bullseye scales along with flower size, meaning the two bullseye domains grow at the same rate. However, the size of the distal domain mostly increases through cell proliferation while the proximal region grows exclusively through cell expansion ^11^ and how this is achieved had remained an unanswered question. Our findings suggest that auxin does not drive the onset of bullseye formation but instead plays a key role controlling cell behaviours in the distal domain. Hence, auxin response is patterned across the petal surface, and this pattern is essential to maintain the bullseye proportions established very early during petal emergence.

First, we found that auxin transcriptional output, as visualized using the transcriptional reporter *DR5v2*, does not correlate with pre-pattern emergence, in both space or time, but instead auxin signalling intensity matches closely with the distribution of cell division events across the petal epidermis: auxin transcriptional output, peaks in the distal and central petal margins from early S0a, when the petal primordium is made of few cells and has just emerged from the inter-sepal zone of the floral meristem. Later, auxin response remains highest in the distalmost epidermal region—and in the corresponding location of presumptive vascular differentiation—however, it is also detected across the epidermis of the entire petal blade, decreasing gradually toward the proximal base. This auxin output pattern mirrors DR5 signals and petal growth profiles previously observed in other species ^44,69,70^. At later stages (S0b to S0c) the pre-pattern is characterised by a medio-lateral asymmetry, with larger cells emerging on the side of the petal most connected to the rest of the floral structure ^11^ and such left-right asymmetry is not detected when the DR5v2 output is recorded. From S0c onward, the high level of auxin signalling in the distal region coincides with regions of high cell proliferation while the DR5v2 response is not clearly detectable in the proximal domain where growth now proceeds via cell expansion instead of division ^11^.

To go beyond correlations, we deployed complementary approaches to manipulate auxin signalling. We used ectopic applications of an auxin transport inhibitor and a scarcely mobile auxin analogue alongside with genetic manipulations *in planta* of transcription factors and enzymes known to modulate auxin distribution and homeostasis. We found that bullseye formation was always robust to those perturbations, with petals successfully establishing contrasting proximal and distal domains in all cases. However, petal growth was impacted due to change in cell proliferation and/or expansion, and the vast majority of growth defect were recorded in the distal bullseye domain. Hence, the distal region of the petal is more sensitive to auxin perturbations and more reliant on fine auxin homeostasis than the proximal domain that forms the bullseye centre. In Arabidopsis, PTL and JAG interface with the auxin signalling pathway at different stages of petal development by organizing local hormone gradients. Changes in the expression of those genes modulate auxin distribution and reduce petal size by affecting polar auxin transport ^32,44,71^. Similarly, constitutive overexpression of *HtPTL1* and *HtJAG1* in Hibiscus results in smaller petals. These phenotypes were largely due to a reduction in the dimensions (length and width) of the distal region, while the length of the proximal domain remains unchanged (but petals are occasionally narrower). We found that reduced cell division accounts for the smaller size of the distal region, consistently with PTL acting as a repressor of cell proliferation in Arabidopsis ^59,60^. In contrast, the shrunken distal domain of *HtJAG1* OE lines was due at least in part to reduced distal cell size. In Arabidopsis, JAG promotes organ outgrowth and cell proliferation in lateral organs, but ectopic or constitutive overexpression can alter how cells coordinate growth with division and yield smaller cells following at least two different routes. First, forcing JAG expression in the floral meristem removes the normal size requirement before genome replication. Consequently, meristematic cells enter S-phase and divide at an abnormally small size ^72^. Second, heterologous overexpression of the tulip tree JAG homolog (*LtuJAG*) reduces *A. thaliana* leaf size by slowing down cell expansion along both the proximo-distal and medio-lateral axes ^73^. Hence, depending on the tissue context, ectopic overexpression of *JAG* can produce smaller cells by overriding the G1/S cell-size checkpoint or by reducing expansion rates and something similar could occur in the developing petal of *HtJAG1* OE *H. trionum* lines.

In addition to affecting cell proliferation, we found that perturbations of auxin distribution also impair conical cell formation as those cells are almost entirely missing in petals treated with high concentrations of 2,4D or petals from *HtGH3.1a OE* lines and *Httcp4.1 Httcp4.2* double mutants. This may result from to two distinct processes. First, auxin signalling could directly contribute to conical cell specification: in *Arabidopsis* petals conical cell expansion correlates with changes in both apoplastic and cytoplasmic pH and with local auxin dynamics ^74^. Hence, non-canonical auxin pathways and rapid auxin responses— involving membrane depolarisation, calcium influx, and cytoskeletal rearrangements—that act independently of the nuclear auxin pathways (NAPs) ^43,75^ could play a role in conical cell elaboration. Fast auxin responses are already present in ancient plant taxa lacking flowers^76^ and future studies with auxin direct sensors^77^, calcium^78^ and pH reporters^79^, and methods for observation of phosphorylation response ^76,80^ are needed to test whether fast auxin responses have been co-opted during flower evolution to contribute to petal patterning. Alternatively, processes at play to maintain organ size could override the cell differentiation process, leading distal cells to expand parallel to the petal surface rather than perpendicularly to try and compensate low cell number by increasing cell size in the petal plane. In that case the change in cell shape morphology observed in the distal region would be a side-effect of mechanisms acting at the tissue scale to maintain organ size, reminiscent of compensation processes originally reported in Drosophila wing and tobacco leaf ^81–83^, rather than be indicative of a direct role for auxin signalling in promoting conical cell formation.

Our work shows that auxin signalling drives distal petal growth via cell proliferation, prompting us to investigate a possible underlying regulatory mechanism. Homologs of TCP4, *HtTCP4.1* and *HtTCP4.2* are preferentially expressed in the distal petal domain as early as S1 and constitutive overexpression of *HtTCP4.1* is sufficient to stimulate cell division in the proximal domains yielding flowers with a larger bullseye ^11^. Hence, we reasoned that TCP4 activity could act upstream of auxin synthesis to promote auxin production and signalling in the distal petal domain, promoting cell proliferation in this region. Our results support this hypothesis as the DR5v2 signal is enhanced in *HtTCP4.1* OE line and transcription of the auxin biosynthetic gene *HtYUCCA3*, whose expression pattern in wild type mirrors the expression of *HtTCP4.1* and *HtTCP4.2*, is enhanced in *HtTCP4.1* OE but reduced in the single *Httcp4.1* mutant. This parallels *Arabidopsis* hypocotyl elongation, where TCP4 directly binds to the promoter of *YUC5*, driving its local expression to produce directional growth ^64^. Similarly, TCP4 interacts with CRABSCLAW and NGATHA transcription factors during gynoecium development to promote the transcription of *YUC2* and *YUC6*, driving localized auxin biosynthesis essential to correct formation of stigmatic papillae ^84^. HtTCP4 could also promote *HtYUC3* expression indirectly: in Arabidopsis TCP4 stimulates the biosynthesis of jasmonic acid by directly activating *LOX2* (*LIPOXYGENASE2*) and elevated jasmonic acid signalling subsequently upregulates *YUC2* and *YUC8*, further increasing endogenous auxin levels ^85^. Notably, *YUC6* overexpression in Arabidopsis does not lead to a uniform DR5 response across hypocotyl tissues ^86^, and likewise, in Hibiscus, *HtTCP4.1* overexpression enhances *HtYUC3* expression across the petal while still maintaining the DR5v2 gradient along the petal proximo-distal axis. This is consistent with the idea that auxin synthesis alone does not account for auxin distribution and patterns of auxin signalling output and suggests the presence of a spatial buffering system in petals likely involving PIN-mediated auxin transport ^29–31^. Interestingly, other TCP genes could contribute to this process as related TCPs (TCP3, GhTCP14) have been shown to impact the expression, polar localization or activity of several PINs and AUX1 proteins central to the auxin transport machinery ^29–32^. What account for the preferential expression of *HtTCP4* homologs in the distal petal domains and whether *HtTCP4* transcription in under control of the pre-patterning processes that divide the petal surface into two territories remains to be established. Assessing whether *HtTCP4* expression is impaired in lines that fail to pre-pattern their petals will be essential to test this hypothesis, but such lines remain to be isolated.

Tissues must be precisely patterned to generate functional organs that allow multicellular organisms to survive and reproduce. This complex process requires the coordination of growth and differentiation. This is particularly important in plants where cells are encased in a wall and laid brick-like. However, we still have a fragmentary understanding of how gene activities drive cell differentiation and growth in floral organs, how the two processes are regulated within and between different subregions of a tissue to ensure correct pattern proportions and how evolution manipulates those mechanisms to generate biodiversity. Petal patterns constitute powerful system to tackle those questions: having established that HtTCP4 activity promotes cell proliferation in the distal domain during *H. trionum* petal development we expected that *Httcp4.1 Httcp4.2* mutants would produce smaller flowers, with a shrunken distal domain but unchanged proximal region. Instead, we obtained flowers similar in size to wild type but with a significantly enlarged bullseye centre. Computational simulations previously indicated that the final bullseye size is determined by the joint regulation of growth and differentiation, and that multiple mechanisms can drive bullseye enlargement ^11^. Our experimental results corroborate this idea, as similarly enlarged bullseyes (21-25% in *HtTCP4.1 OE* and *Httcp4.1-1,Httcp4.2-6* vs. 14.5-15% in wild type) stem from distinct changes in cell proliferation or expansion in the different bullseye domains. This also highlights the existence of compensatory mechanisms, where conical cells are replaced by flat elongated tabular cell that almost offset the 31% reduction in cell number in the distal region of the double mutant. Such mechanisms at work to maintain petal size at the organ level are poorly understood and make it difficult to predict the impact of modifications affecting molecular regulators of cell behaviour. Future work aiming at deciphering the GRNs controlling growth through the different regions of the petal and their interplay with the signalling and mechanical processes involved in petal size maintenance will be essential to develop predictive models to modulate bullseye proportions at will.

Bullseye size is ecologically relevant and can impact flowering plant fitness and speciation ^6,58,87^. Here we uncovered a role for auxin signalling in maintaining bullseye proportions (by controlling differential growth across the petal during bullseye formation) while the petal size increases dramatically. We found that chemical or genetic perturbations disrupting the auxin output map often led to a change in bullseye dimensions hence auxin signalling could act as a pattern modifier ^9,10^. It is tempting to speculate that mutations affecting auxin synthesis, homeostasis, transport or signalling may have contributed to the emergence of bullseye size variants during evolution, but this remains to be tested. Future efforts to sequence collections of novel ecotypes and map the genetic bases of natural phenotypic variation across diverse species should provide valuable data to test this hypothesis and more generally, assess the contribution of phytohormones to the evolution of the vast morphological diversity of flowering plants.

## MATERIALS AND METHODS

### Plant material and growth condition

Wild-type *H. trionum L*. CUBG seeds (voucher CGE00046422) were originally obtained from Cambridge University Botanic Gardens, Cambridge, UK and subsequently maintained as stock at the Sainsbury Laboratory Cambridge University (SLCU). The *HtTCP4.1* OE (overexpression) line was originally published in ^11^. All other plant lines were generated for this study. All lines used in this study are listed in **Supplementary Data 1**. Seeds were imbibed in 90°C water for 10 minutes and germinated in the dark at 30°C for 48h. Seedling were transferred to a mix of 1:3 parts of Levington’s (UK) M3 and Advance F2 seed & modular composts and grown in the glasshouse facilities at the Sainsbury Laboratory (University of Cambridge, UK) under controlled conditions on a 16h:8h, light:dark photoperiod at 24°C with a minimum radiance of 88 W.m^-2^.

### Ectopic 2*,4-D* and *TIBA* treatments

Healthy and similarly developed plants were selected for treatment and assigned either to the treatment containing the active principle or the mock solution. Floral buds were treated daily beginning before stage 1 (unpatterned petals) and continuing through anthesis (stage 5). Treatments were made with auxin 2,4-dichlorophenoxyacetic acid (2,4-D) (SIGMA-Aldrich) and the auxin transport inhibitor 2,3,5-triiodobenzoic acid (TIBA) (SIGMA-Aldrich) dissolved in dimethyl sulfoxide (DMSO) and ethanol and then diluted with distilled water and 0.01% Silwet L-77 solution to the desired working concentrations (1 µM, 2.5 µM, 5, µM 10 µM, 50 µM, 100 µM for 2,4-D; 1 µM, 10 µM, 50 µM, 100 µM for TIBA). The mock solutions used to treat the control plants only included the solvent solutions appropriately diluted plus 0.01% Silwet L-77 without active compounds. The final solution contained 0.01% (v/v) Silwet L-77 (Lehle Seeds, USA) as a surfactant to enhance penetration and adhesion on samples.

### Construction of plant expression vectors for gene overexpression and *DR5v2* reporter construction

All the gene coding sequences were retrieved from the genome assembly Htrionum_1.0 (Submitted GenBank assembly GCA_030270665.1) ^57^ and the used gene IDs are listed in **Supplementary Data 2**.

The complete *DR5v2::ntdTomato::NOSt* cassette was Eco RI (New England Biolabs)-digested and cut from the original plasmid *DR5v2* in pGreenIIK containing the 7x(TGTCGG) auxin responsive element (*DR5v2*) driving the expression of the fluorescent protein *ntdTomato*, followed by the nopaline synthase terminator described by ^52^ and kindly provided by Pau Formosa-Jordan and Dolf Weijers. The cut cassette was assembled into EcoRI-linearized pSG26 vector using NEB DNA Ligase (New England Biolab). pSG26 is a modified pCAMBIA binary vector carrying a *pUBQ10::eYFP::NOSt* transformation-marker cassette and a plant kanamycin-resistance cassette.

The *TCSnew::tdTomato–DR5v2::GFP* dual-reporter region was amplified by Q5 High-Fidelity DNA Polymerase (New England Biolabs) PCR from the construct donated by Prof. Bert De Rybel described by ^88^ and assembled into the linearized pSG146 binary vector (modified pCAMBIA binary vector containing a pUBQ10::AtPIP2A–mTurquoise2 and a plant kanamycin-resistance cassette) using NEBuilder HiFi DNA Assembly (New England Biolabs). The resulting construct retained the *pUBQ10::AtPIP2A–mTurquoise2* cassette.

The coding sequences (cds) of *HtJAG1*, *HtPTL1*, *HtGH3.1a*, *HtGH3.1 b*, *HtGH3.1c*, and *HtIAMT1* were amplified using Q5 High-Fidelity DNA Polymerase with the gene-specific primers listed in **Supplementary Data 4**. PCR products were initially subcloned into EcoRV-linearized pBluescript KS(−) for intermediate propagation. The coding sequences were subsequently excised from the intermediate constructs using the restriction enzymes listed in **Supplementary Data 3**. The *HtGH3.1a* coding sequence was inserted downstream of the CaMV 2×35S promoter in a modified pGreenII binary vector. The remaining coding sequences were cloned downstream of the CaMV 2×35S promoter in modified pCAMBIA binary vectors. An additional HtGH3.1-like a overexpression construct was generated using the Arabidopsis UBIQUITIN10 promoter. The modified pGreenII and pCAMBIA vectors carried a pUBQ10::eYFP cassette, allowing identification of transformed plants through eYFP fluorescence in planta.

All final constructs were verified by Sanger sequencing (Azenta Life Sciences) before plant transformation. A summary of all final constructs used for plant transformation is provided in **Supplementary Data 3**.

### Generation of CRISPR/Cas9 vectors for gene editing

Oligonucleotides (Sigma-Aldrich) used to generate all the spacer targeting genes of interest (HtTCP4.1 and HtTCP4.2) flanked by PAM sequences in the *H. trionum* genome were designed feeding the web tool https://eu.idtdna.com/site/order/designtool/index/CRISPR_SEQUENCE with the genomic sequence of the GOI and were designed to generate BstI restriction sites upon digestion. Oligonucleotide sequences and the corresponding target GOIs are given in **Supplementary Data 4**.

To assemble each guideRNA coding sequence, oligonucleotides were annealed to generate double strand spacers that were then ligated into BstI-digested pSG23 (pBSK:pAtU6:26-guideRNA) using T4 DNA ligase (New England Biolabs). Ligated constructs were transformed into *E. coli* DH5α cells and plated on carbenicillin-containing LB medium. Two distinct pSG23-sgRNA vectors (level 1 vector) were generated to target each GOIs. Positive clones were verified by colony PCR and Sanger sequencing (Genewiz). The Level 1 vectors were used as templates to amplify the guideRNAs sequences using Q5 High-Fidelity DNA Polymerase (New England Biolabs) and generate inserts compatible with the downstream application of the NEBuilder assembly system. The final plant expression vector (level 2) was produces using the NEBuilder assembly system (New England Biolabs) following manufacturer’s instruction to combine the two guidRNA sequences targeting each GOI with the pSG100 binary vector (pBSK:pUBQ10-pcoCAS9-UBQ10-eYFPmb) that contains two AtU6 promoters for sgRNA expression, a Cas9 coding sequence, a YFPmp constitutively expressed under control of AtUBQ10 promoter (serve as marker to quickly identify transformants) and an nptII gene for kanamycin selection. NEBuilder reaction products were transformed into *E. coli* DH5α cells and plated on kanamycin-containing LB medium. Positive clones were verified by colony PCR and Sanger sequencing (Genewiz). A summary of all final CRISPR/Cas9 constructs used for plant transformation is provided in **Supplementary Data 3**.

### Plant transformation

Electrocompetent *Agrobacterium tumefaciens* LBA4404 cells were transformed with plant expression vectors described in the corresponding section. Transformed cells were selected with 50 mg/L kanamycin and 25 mg/L streptomycin. Transgenic *H. trionum* lines were produced using agrobacterium-mediated transgene delivery followed by tissue culture to induce callus production and plant regeneration, as detailed in ^58^. Positive transformants regenerated plants were selected under a fluorescent stereoscope (Zeiss Lumar V12) for visualizing YFP (Yellow Fluorescent Protein) using a YFP longpass filter.

### Screening for genome edited lines

Regenerated plantlets (T0) were initially screened for transgene expression using a Zeiss Lumar V12 fluorescent stereomicroscope to identify eYFPmb-positive individuals. Genomic DNA was then extracted from fluorescent plants using a modified alkaline lysis protocol. Briefly, leaf tissues were homogenized in lysis buffer (100 mM Tris-HCl pH 8.0, 250 mM KCl, 10 mM EDTA; final pH adjusted to 9.3), followed by incubation at 65 °C for 10 minutes. After lysis, samples were diluted with 3% BSA solution (pH 7.6) and further diluted in nuclease-free water for downstream PCR applications.

PCR-based genotyping was performed using primers flanking the targeted genomic loci of the genes of interest (GOIs), followed by Sanger sequencing using appropriate primers to confirm editing events in T0 lines. The list of primers used for genotyping is provided in **Supplementary Data 4**.

Confirmed T0 edited plants were allowed to self-pollinate, and T1 seeds were pre-screened using the Zeiss Lumar V12 stereomicroscope to identify non-fluorescent individuals, indicating segregation of the Cas9-mYFPmb transgene. Genomic DNA was extracted from these individuals, and the target GOIs were amplified by PCR and validated by Sanger sequencing.

T2 segregating lines lacking YFP fluorescence were further analyzed to confirm homozygous editing and absence of the Cas9-mYFPmb transgenes. Only confirmed homozygous, transgene-free plants were used for phenotypic analyses and their alleles are detailed in **Supplementary Data 5**.

### Petal phenotyping and image analysis

Flowers at anthesis (S5) were collected in the morning and imaged on black velvet using a Panasonic DMC-FZ38 camera (Panasonic Corporation). Dissected petals were imaged with a Keyence microscope (VHX7000) fitted with VH-Z100R lens at magnifications of 20X, 150X, and 700X for the cell size analyses. Petals with the adaxial side facing upward were scanned (Epson Perfection V850 Pro) on black and white backgrounds. Petal white and purple pigmented sizes and petal lengths were measured from scanned images. All petal features were manually measured using ImageJ (v.1.53q https://imagej.net/ij/). For each flower, 4-5 petals or 10 cells were measured and averaged from a minimum of 10 flowers per genotype/ transformant line. Statistical analyses were performed using RStudio (Version 1.1.1717).

All statistical analyses were conducted using RStudio (version 1.1.1717). Prior to hypothesis testing, data were assessed for normality using the Shapiro–Wilk test and for homogeneity of variance using Levene’s test. Depending on the distribution and variance characteristics of the data: if assumptions of normality and equal variance were met, standard t-tests (for two-group comparisons) or one-way ANOVA (for multiple groups) were used. When variances were unequal, but normality was respected, Welch’s t-test or Welch’s ANOVA was applied. For data that did not meet normality or variance assumptions, non-parametric tests such as the Mann–Whitney U test (two groups) or the Kruskal–Wallis test (multiple groups) were used. For post hoc analysis following significant ANOVA, the appropriate pairwise comparisons were conducted using Student’s t-test, Welch’s t-test, or Wilcoxon rank-sum test, based on the underlying distribution and variance structure of the data.

### Analysis of the distribution of the cell geometry features (cell area) and *DR5v2* signal across the petal primordia using confocal microscopy

For all confocal imaging experiments, petal primordia were dissected with fine tweezers from floral buds at developmental stages from S0a to S1, as defined in ^11^. Dissected petals were mounted adaxial side up on Petri dishes using double-sided adhesive tape under a Zeiss stereomicroscope. To reconstruct auxin response maps, petals expressing *DR5v2::ntdTomato* or *DR5v2::GFP* were mounted in water and imaged using either a ZEISS LSM 880 confocal microscope with a 20× water-dipping Apochromat objective, or a Leica SP8 confocal microscope with a 25× water-dipping Apochromat objective using the LasX Navigator software for automated tiling and Z-stack acquisition. Fluorescent reporters were excited and detected using the following laser lines and emission filter settings:

– 561 nm for ntdTomato (emission: 580–630 nm);
– 488 nm for eGFP (emission: 520–560 nm);
– 514 nm for eYFP (emission: 500–550 nm).

Imaging parameters (laser power, main and digital gain, digital offset) were kept constant. Z-stacks were acquired to cover the entire petal surface, and images were stitched using Imaris Stitcher (Oxford Instruments). Fluorescence quantification was performed using Imaris (v9.3, Oxford Instruments), MorphoGraphX (v2.0) or ImageJ (v1.53q, for the CTCF for the *DR5v2* responsivity to auxin treatments in roots).

### Nuclear Fluorescence Quantification in Imaris and data analysis

Nuclear segmentation and *DR5v2* fluorescence measurements were performed using Imaris v9.3 (Oxford Instruments). The following pipeline was used for each 3D confocal image stack: (i) *Spot* model based on the measured average nuclear diameter (∼6–8 μm) to carry out the nuclear segmentation, (ii) *Background subtraction* to enhance nuclear signal-to-noise ratio, (iii) *Automatic quality thresholding* on each sample to exclude false-positive detections, and (iv) Manual corrections of the nuclear segmentation to exclude spurious signal.

For each detected nucleus, mean fluorescent intensity within the spot volume was calculated from the *DR5v2* channel. Only nuclei with fluorescence intensities above the background threshold were considered DR5v2-positive and included in downstream analyses. The (x, y, z) spatial coordinates of each nucleus relative to the manually positioned base of the petal (origin of the axes) were also extracted to assess positional information along the proximo-distal (PD) axis scaled between 0 (base) and 1 (tip). Across samples, the same spot detection parameters were applied within each stage and replicate. For S0a, entire petals were analysed. For S0b, S0c, and S1, only a central longitudinal stripe (∼20% of the maximum petal width) was extracted to minimize lateral positional noise. ROIs were manually drawn and applied consistently across replicates.

To analyse the distribution of *DR5v2* signal along the petal proximo-distal (P-D) axis, generalised additive mixed models (GAMMs) were applied using the mgcv package in R (R version 4.4.2). The analysis was performed following the steps below.

*Data Processing:*

DR5v2 intensity values were normalized per sample (Relative_Intensity = Intensity_n_/Intensity_MAX_)

The *y* component of the PD coordinate was scaled between 0 (petal base) and 1 (tip) (Relative_Position = _Position = *y_i_*/*L*) where L is the maximum length from the base to the tip of the petal.

Rolling averages of normalised fluorescence intensity were computed within each sample using the zoo::rollapply() function, with a window size of 10 measurements, to reduce noise prior to model fitting.

*Model Specification:*

-*Fixed effects:* position along the PD axis ((Relative_Position = *y_i_*/*L*) modelled a smooth term to capture the shape of the signal profile;

*-Random effects:* a random intercept for sample (individual petal) to account for measurements grouped within petals. biological replicate and individual petal identity to account for nested data structure and repeated measurements,

*-Response variable:* the rolling average of normalized *DR5v2* fluorescence intensity ((Relative_Intensity = Intensityn/IntensityMAX).

Models were fitted separately for each developmental stage using the gamm() function from the mgcv package, with Gaussian errors and an identity link. Model fits were visualised as DR5v2 distributions along the PD axis using model-based smooth curves (red), generated from the GAM component of each fitted model using predict(…, type = “response”, se.fit = TRUE). Curves were overlaid on the rolling-average intensity measurements using ggplot2::geom_line(), and approximate pointwise 95% confidence intervals were displayed using ggplot2::geom_ribbon(), calculated as the fitted values ± 1.96 standard errors.

### Cell area Quantification and *DR5v2* fluorescent quantification and mapping using MorphoGraphX

Petals were stained with membrane-specific dyes according to the following conditions:

-0.01 μg/μL FM4-64 (Thermo Fisher Scientific) was used to stain membranes for simultaneous analysis of *DR5v2* signal and cell geometry in *DR5v2:ntdTomato* lines at stages S0a and S0b, with incubation times ranging from 5 to 20 minutes.

Stained petals were rinsed twice in distilled water prior to imaging. FM4-64 was excited with a 514 nm laser line, and fluorescence was collected between 537–622 nm. Z-stacks were acquired at a 2.5 μm step size to cover the full petal surface using the LasX Navigator module on the Leica SP8.

Image segmentation was performed using MorphoGraphX (v2.0). Cell membranes were segmented using the “auto-seed” and “cell segmentation” tools adapting the following parameters: *membrane signal threshold*: 40–60 (adjusted per image for optimal edge detection), *Seed smoothing*: 2–3 and *Watershed sensitivity*: medium

All segmentation outputs were manually corrected to fix boundary errors. Final 2.5D cell meshes were flattened into 2D meshes, and the *DR5v2* signal intensity was projected onto the segmented mesh surface from the relevant confocal channel.

Quantitative data on spatial coordinates, on cell geometry (area) at S0a and S0b and data on *DR5v2* fluorescence intensity at S0a and b of *DR5v2:ntdtomato* were exported as csv files and processed using RStudio (v1.1.1717). Data analysis was performed using the following R packages: tidyverse, ggplot2, rstatix, reshape2, ggpubr. The *DR5v2* data processing, model specification and visualization for statistical analysis was conducted using the pipeline outlined above in “Nuclear Fluorescence Quantification in Imaris and data statistical modelling”.

### Statistical Correlation Analyses Between Auxin Response and Cell Size at Stage 0a and Stage 0b

Correlation analyses between relative *DR5v2* fluorescent intensity and relative cell area at S0a and S0b were conducted in RStudio using built-in and specialized statistical packages. Associations between relative *DR5v2::ntdTomato* fluorescence intensity (Iᵢ/I_max_) and relative cell area (Aᵢ/A_max_) were analysed separately at stages S0a and S0b in R. Pearson’s correlation coefficient was calculated from paired measurements within each sample, (n=5 per stage). One overall estimate per stage was computed accounting for measurements grouped within samples through Fisher z-transformation of sample-level correlation coefficients and averaging with equal weight assigned to each sample. Mean correlations and their 95% confidence intervals were obtained by back-transforming the mean Fisher z values and their Student’s t-based confidence limits. Two-sided one-sample t-tests assessed whether the mean Fisher z differed from zero (df = 4). P values were adjusted using the Holm procedure across the two stage-specific tests.

### Tissue collection & RNA extraction

Total RNA extraction was performed on Stage 3 petals from CUBG wildtype *H. trionum*, *HtTCP4.1* OE and *Httcp4.1-2* mutant. Three biological replicates were collected for each stage and genotype. Frozen petal tissues were ground to a fine powder and total RNA was extracted using the Spectrum Plant Total RNA kit (Sigma STRN250) with ∼40 mg of ground tissue as starting material for each RNA extraction.

### cDNA synthesis

cDNA was synthesized from 0.5 ug total RNA using SuperScript III Reverse Transcriptase (Invitrogen) following manufacturer’s guidelines. Oligo(dT)15 primers were used for first strand cDNA synthesis for gene isolation and measuring expression by RT-PCR or qPCR. First-strand cDNA synthesis followed manufacturer’s guidelines, with incubation with SuperScript III RT at 50°C for 1 hr followed by an inactivation step at 70°C for 15 min.

### Verification of transgene overexpression using RT-PCR

Transgene overexpression was verified by semi-quantitative reverse-transcription PCR (semi-qRT-PCR) in independent transgenic lines and compared with wild-type H. trionum CUBG. Total RNA was extracted from leaf tissue and reverse-transcribed into cDNA. Gene-specific amplification was performed for 30 PCR cycles, using primers listed in **Supplementary Data 4**, with *HtACTIN1* used as a reference gene. PCR products were separated by agarose gel electrophoresis, and increased transcript accumulation in the overexpression lines relative to wild type was assessed from band intensity. Electrophoresis gels confirming transgene overexpression are shown in **Supplementary Fig. 3, 4**.

### Analysis of gene expression level using Quantitative Real-Time PC

Quantitative RT-PCR was performed to quantify expression levels of the GOIs using the specified oligonucleotides in **Supplementary Data 4 (**target gene IDs listed in **Supplementary Data 2**) using a Roche LightCycler 480 machine and the Luna Universal qPCR Master Mix (New England BioLabs). *HtACTIN1* was used as housekeeper gene for all experiments as in ^2^. Data was analyzed using an efficiency-corrected ΔΔCt and the *HtACTIN1* reference gene for normalization ^89,90^. Technical-replicate Ct values were averaged separately for the target and reference genes within each biological replicate. Efficiency-corrected ΔCt was calculated as Ct_target × log₂(E_target) − Ct_reference × log₂(E_reference). Within each experiment and petal region, the mean corrected ΔCt of the CUBG biological replicates served as the common calibrator. Log₂ relative expression was calculated as −ΔΔCt. Statistical analyses were calculated in R using RStudio (Version 1.1.1717 Comparisons between each transgenic genotype and its corresponding CUBG control were performed separately for proximal and distal regions using two-sided Welch’s t-tests on log₂ relative expression values. P-values were adjusted using the Benjamini–Hochberg procedure across the four planned comparisons. Each plotted point represents one biological replicate, and horizontal bars indicate group means.

## Acknowledgments

The authors thank Ray Wightman and Gareth Evans for their help with microscopy; SLCU horticulture team for plant care; Marco Palma (University College London) for discussing statistical modelling and suggestions on R analyses; Selin Zeyrek for productive discussions of the results.

## Author Contributions

Design of the project: E.M. Conceptualization: E.M. and E.S. Data curation: E.S., S.G., M.T.S.Y. and L.R. Formal analysis: E.S. Funding acquisition: E.M., E.S. Investigation: E.S., S.G., E.M., M.T.S.Y., L.R., and E.D. Methodology: E.M., E.S., S.G., M.T.S.Y., L.R, E.D. Resources: E.M., E.S., S.G., M.T.S.Y. Supervision: E.M. Validation: E.M., E.S. Visualization: E.S. and E.M. Writing—original draft: E.S. and E.M. Writing—review and editing: E.S., E.M, M.T.S.Y., L.R and E.D.

