## Supplementary Figures and Legends for "Auxin regulates the proportions, but not the initiation, of *Hibiscus trionum* bullseyes"

**Article title**

The following Supporting Information is available for this article:

**Supplementary Figure 1**
**Supplementary Figure 2**
**Supplementary Figure 3**
**Supplementary Figure 4**
**Supplementary Figure 5**

The following five Supplementary Data files are provided as separate files:

**Supplementary Data 1**. *H. trionum* transgenic and genome edited lines generated in this study.
**Supplementary Data 2**. Gene identifiers.
**Supplementary Data 3**. Plant expression vectors generated for this study.
**Supplementary Data 4**. Primer sequences used in this study.
**Supplementary Data 5**. Mutated alleles for *HtTCP4.1* and *HtTCP4.2* generated with CRISPR/Cas9 genome editing for this study

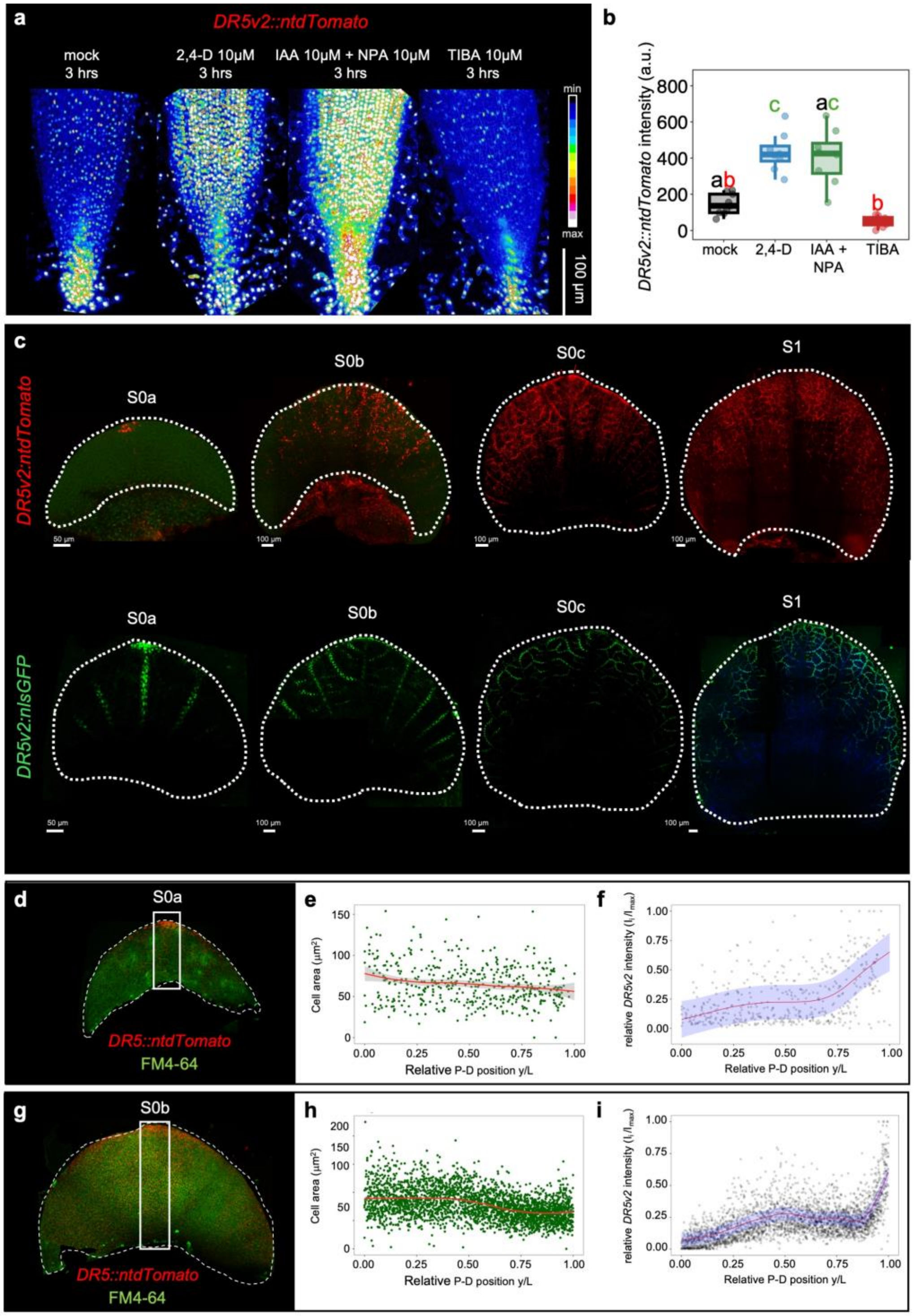


**Supplementary Figure 1.** (**a**) Representative Maximum Intensity Projections of LCSM images of *DR5v2::ntdTomato* root tips following mock, 10 µM 2,4-D, 10 µM IAA + 10 µM NPA, or 10 µM TIBA treatments for 3 hours. *DR5v2::ntdTomato* is false-coloured according to the heatmap shown on the right. (**b**) *DR5v2::ntdTomato* fluorescence intensity following mock, 10 µM 2,4-D, 10 µM IAA + 10 µM NPA, or 10 µM TIBA treatment for 3 hrs. Compared with mock (144.6 ± 58.2 a.u.), mean fluorescence intensity was increased by 199% following 2,4-D treatment (432.6 ± 107.8 a.u.; adjusted P = 0.0464) and by 181%

following IAA + NPA treatment (406.3 ± 152.9 a.u.; adjusted P = 0.0737), and reduced by 66% following TIBA treatment (49.8 ± 30.8 a.u.; adjusted P = 0.478). Values are mean ± SD. Each point represents one quantified sample; n = 9 for mock and n = 8 for each treatment. Overall differences assessed with Kruskal–Wallis test [H(3) = 25.71, P = $1.10 \times 10^{-5}$]. Pairwise comparisons used two-sided Dunn tests with Bonferroni correction across all comparisons. Groups sharing at least one letter were not significantly different; groups sharing no letters differed at adjusted P < 0.05.

(**c**) Representative maximum intensity projections of LCSM images of *DR5v2::ntdTomato* (top row) and *DR5v2::nlsGFP* (bottom row) petal primordia at S0a-S1 as indicated for comparison.

(**d**) Representative maximum intensity projection LCSM image of *DR5v2::ntdTomato* petal primordium stained with FM4-64 at stage S0a. (**e**) Distribution plot of epidermal cell areas ($\mu m^2$) along the normalized proximodistal axis ($y_i/L$) of S0a petals. Points represent individual cell measurements, the red line shows a locally estimated scatterplot smoothing (LOESS) fit, and shading indicates the approximate pointwise 95% confidence interval around the fitted profile. (**f**) Distribution of *DR5v2::ntdTomato* relative fluorescence intensity ($I_i/I_{max}$) along the normalized proximodistal axis ($y_i/L$) of S0a petals. Points represent rolling-average fluorescence intensities calculated within each sample using a window of 10 measurements.The red line shows the mean profile estimated using a generalized additive mixed model (GAMM), with a smooth effect of relative position and a random intercept for sample. Blue shading indicates the approximate pointwise 95% confidence interval around the fitted mean profile (estimate ± 1.96 standard errors).
For e-f, 453 paired-measurements, from n=5.

(**g**) Representative maximum intensity projections LCSM images of *DR5v2::ntdTomato* petal primordium stained with FM4-64 at stage S0b. (**h**) Distribution plot of epidermal cell areas ($\mu m^2$) along the normalized proximodistal axis ($y_i/L$) of S0b petals. Points represent individual cell measurements, the red line shows a locally estimated scatterplot smoothing (LOESS) fit, and shading indicates the approximate pointwise 95% confidence interval around the fitted profile. (**i**) Distribution of *DR5v2::ntdTomato* relative fluorescence intensity ($I_i/I_{max}$) along the normalized proximodistal axis ($y_i/L$) of S0b petals. Points represent individual measurements. The red line shows the mean profile estimated using a generalized additive mixed model (GAMM), with a smooth effect of relative position and a random intercept for sample. Blue shading indicates the approximate pointwise 95% confidence interval around the fitted mean profile (estimate ± 1.96 standard errors).
For h-I, 2852 paired-measurements, from n=5.

**a**

| Reduction (%) = 100 × (1 − mean treatment / mean mock) | | | |
|---|---|---|---|
| TIBA [ ] | Proximal reduction (% relative to mock) | Distal reduction (% relative to mock) | Total length reduction (% relative to mock) |
| 1 µM | 0.2% | 9.3% | 6.5% |
| 10 µM | 3.7% | 7.7% | 6.4% |
| 50 µM | 16.8% | 19.6% | 18.8% |
| 100 µM | 22.5% | 26.9% | 25.5% |

**b**

| Reduction (%) = 100 × (1 − mean treatment / mean mock) | | | |
|---|---|---|---|
| 2,4-D [ ] | Proximal reduction (% relative to mock) | Distal reduction (% relative to mock) | Total length reduction (% relative to mock) |
| 2.5 µM | 10.0% | 44.2% | 32.9% |
| 5 µM | 1.9% | 49.8% | 33.9% |
| 10 µM | 24.8% | 59.6% | 48.0% |
| 50 µM | 31.3% | 62.6% | 52.2% |
| 100 µM | 38.3% | 60.8% | 56.6% |

**Supplementary Figure 2.** (**a**) Summary of percentage changes relative to mock for all TIBA treatment concentrations (µM). (**b**) Summary of percentage changes relative to mock for all 2,4-D treatment concentrations (µM).

.

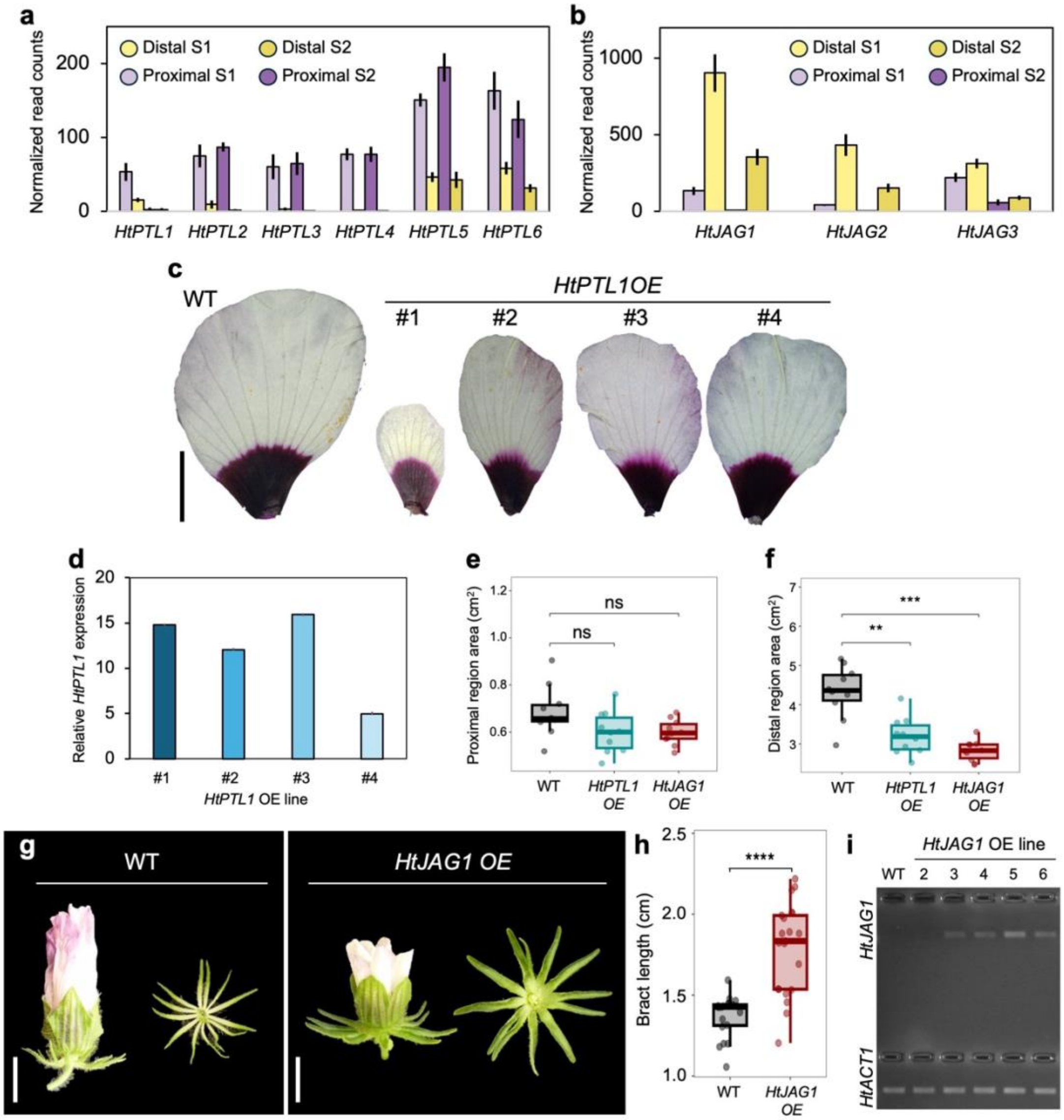


**Supplementary Figure 3.** Normalized read counts for (**a**) the six *PTL* homologs and (**b**) the three *JAG* homologs in *H. trionum* from bulk RNA-seq of proximal and distal dissected stage 1 (respectively P1 and D1) petals and stage 2 (respectively P2 and D2) petals. Information extracted from original datasets published in Yeo et al., 2025 and Oud et al., 2026. (**c**) Representative *Hibiscus trionum* petals at anthesis (S5; right) from wild type and *HtPTL*1 OE lines #1, #2, #3 and #4. Scalebar = 1cm. (**d**) qRT-PCR quantification of *HtPTL1* relative transcript abundance (fold change as $\log_2$ expression relative to wild type) in petal tissue of *HtPTL*1 OE lines #1, #2, #3 and #4. (**e**) Area of proximal region ($cm^2$) of S5 petal from wild type, *HtPTL1* OE and *HtJAG1* OE lines. Differences in mean area were not statistically significant ($0.600 \pm 0.088$ cm² in *HtPTL1* OE vs. $0.686 \pm 0.107$ cm² in wild type; two-sided Welch's t-test, t = 1.964, df = 17.38, Holm-adjusted P = 0.0772 and $0.600 \pm 0.053$ cm² in *HtJAG1* OE vs. $0.686 \pm 0.107$ cm²; t = 2.297, df = 13.21, Holm-adjusted P = 0.0772). (**f**) Area of distal region ($cm^2$) of S5 petals from wild type, *HtPTL1* OE and *HtJAG1* OE lines. Mean distal region area was reduced by 26% in *HtPTL1* OE ($3.20 \pm 0.47$ cm² vs. $4.33 \pm 0.67$ cm²; t = 4.344, df = 16.16, Holm-adjusted P = 0.00148) and by 35% in *HtJAG1* OE ($2.83 \pm 0.26$ versus $4.33 \pm 0.67$ cm²; t = 6.612, df = 11.62, Holm-adjusted P = 0.000117).
For panels e-f, values are mean ± SD; percentage changes are relative to the corresponding wild type mean. Each point represents one quantified sample; n= 10 per genotype. Two-sided Welch's *t*-tests compared each

OE line with WT within each region, with Holm correction across comparisons. ns = not significant (adjusted p-value $\geq 0.05$), ** = adjusted p-value $< 0.01$, *** = adjusted p-value $< 0.001$.

(**g**) Representative *Hibiscus trionum* flower just after anthesis (left, side view) and associated whorl of bracts (right, bottom view) from wild type and *HtJAG*1 *OE* line #5. Scalebar = 1cm. (**h**) Bract length (cm) in wild type and *HtJAG*1 *OE* line #5. Mean bract length was increased by 30.1% in the OE line compared with WT (1.786 ± 0.279 versus 1.373 ± 0.128 cm; means ± SD; two-sided Welch's t-test, $t = -6.144$, $df = 28.30$, $P = 1.19 \times 10^{-6}$). Values are mean ± SD, and percentage changes are calculated relative to the corresponding wild type mean. $n = 20$ flowers for WT and $n = 20$ for the *HtJAG*1 *OE* line #5. In the plot, each point represents one flower. **** = p-value $< 0.0001$. (*i*) Semi-quantitative RT-PCR analysis of *HtJAG1* expression in *HtJAG1* OE lines #2, #3, #4, #5, #6 and wild type (CUBG). Gel electrophoresis shows amplification products obtained from cDNA from leaf tissue using primers for *HtJAG1* gene (top) and the reference gene *HtACT1* (bottom), after 30 PCR cycles.

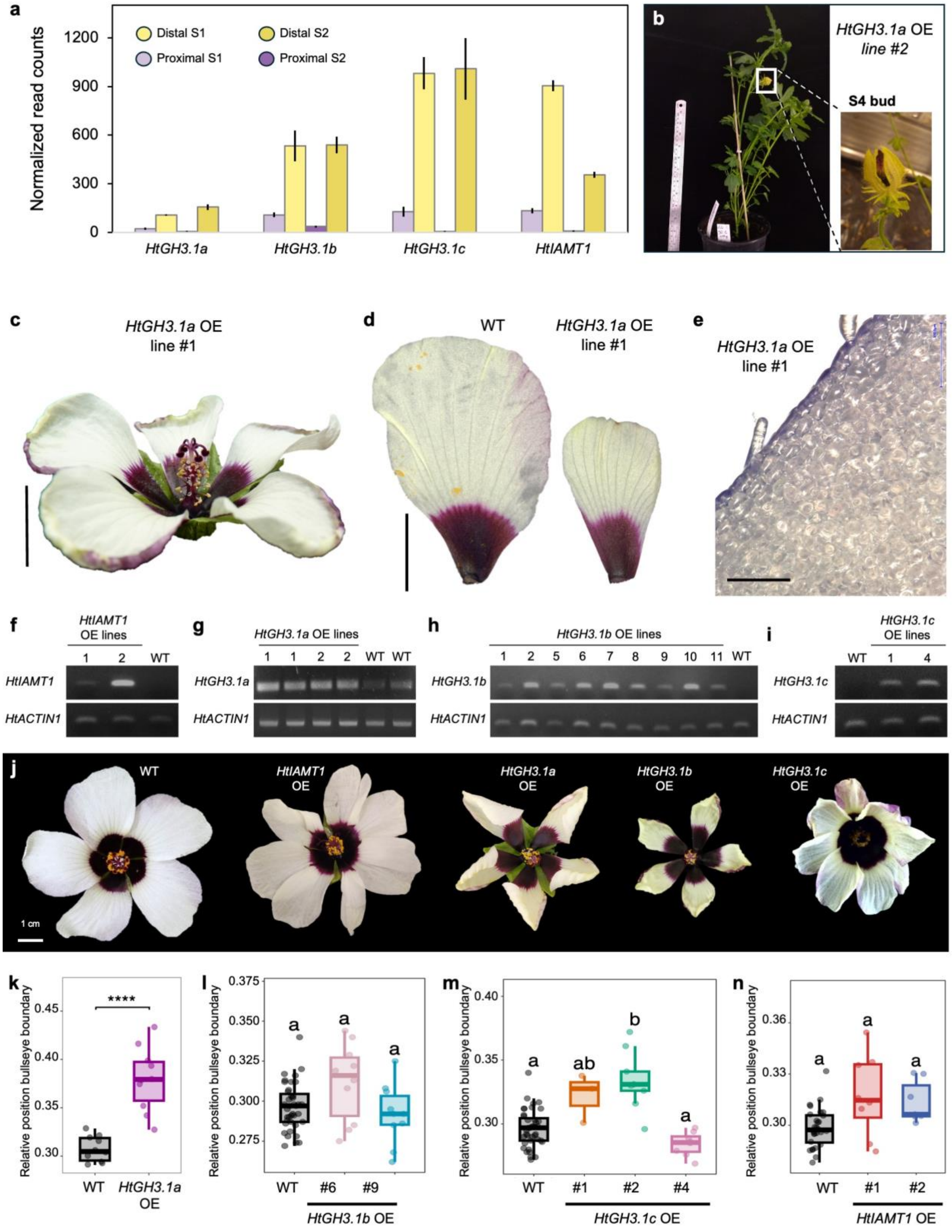


**Supplementary Figure 4.** (**a**) Normalized read counts for *HtGH3.1a, HtGH3.1b*, *HtGH3.1c* and *HtIAMT1* from bulk RNA-seq of proximal and distal dissected stage 1 (respectively P1 and D1) petals and stage 2

(respectively P2 and D2) petals. Information extracted from original datasets published in Yeo et al., 2025 and Oud et al., 2026. (**b**) *HtGH3.1a* OE line #2 showing dead buds failing to reach anthesis (S5) on the left and a close-up of dead S4 bud on the right. (**c**) Flowers of *HtGH3.1a* OE line #1 at S5 seen from a lateral view, showing petals curling up at borders. Scalebar = 1 cm. (**d**) Petals of wild type and *HtGH3.1a* OE line #1 at S5, showing reduced petal size but comparable proximal region length. Scale bar = 1 cm. (**e**) Close-up of the distalmost region of *HtGH3.1a* OE line #1 petals at S5, showing flattened cells in place of conical cells. (**f**) Semi-quantitative RT-PCR analysis of *HtIAMT1* expression in *HtIAMT1* OE line #1 and line #2, and wild type (CUBG). Gel electrophoresis shows amplification products obtained from cDNA using primers for *HtIAMT1* (top) and the reference gene *HtACT1* (bottom), after 30 PCR cycles. (**g**) Semi-quantitative RT-PCR analysis of *HtGH3.1a* expression in *HtGH3.1a* OE lines *#1* and *#2* and wild type (CUBG). Gel electrophoresis shows amplification products obtained from cDNA using primers for *HtGH3.1a* (top) and the reference gene *HtACT1* (bottom), after 30 PCR cycles. (**h**) Semi-quantitative RT-PCR analysis of *HtGH3.1b* expression in *HtGH3.1b* OE lines #1, #2, #5, #6, #7, #8, #9, #10, and #11, and wild type (CUBG). Gel electrophoresis shows amplification products obtained from cDNA using primers for *HtGH3.1b* (top) and the reference gene *HtACT1* (bottom), after 30 PCR cycles. (**i**) Semi-quantitative RT-PCR analysis of *HtGH3.1c* expression in *HtGH3.1c* OE line #1 and line #4, and wild type (CUBG). Gel electrophoresis shows amplification products obtained from cDNA using primers for *HtGH3.1c* (top) and the reference gene *HtACT1* (bottom), following 30 PCR cycles. No cDNA could be retrieved for line #2. (**j**) Additional images of *Hibiscus trionum* flowers of wild type (CUBG), *HtIAMT1* OE*, HtGH3.1a* OE*, HtGH3.1b* OE*,* and *HtGH3.1c* OE at S5. Scalebars = 1 cm.

(**k**) Relative position of the bullseye boundary along the petal proximo-distal axis in wild type and *HtGH3.1a* OE line #1, calculated as the ratio between the length of the proximal region length and the total petal length. The OE line showed a 23% increase in mean bullseye proportion relative to wild type (0.38 ± 0.03 versus 0.31 ± 0.01; mean ± SD, two-sided Welch's t-test, t {11.894} = -6.440, P = 3.344 $10^{-5}$).

(**l**) Relative position of the bullseye boundary along the petal proximo-distal axis in wild type and *HtGH3.1b* OE line #6 and line #9, calculated as the ratio between the length of the proximal region length and the total petal length. Differences were not statistically significant (Holm-adjusted P=1.00 for both lines).

(**m**) Relative position of the bullseye boundary along the petal proximo-distal axis in wild type and *HtGH3.1c* OE lines #1, 2 and 4, calculated as the ratio between the length of the proximal region length and the total petal length. The relative position of the boundary only significantly differs from wild type in line #2 (12.60% higher in line #2, Holm-adjusted P=4.55 x$10^{-4}$).

(**n**) Relative position of the bullseye boundary along the petal proximo-distal axis in wild type and *HtIAMT1* OE #1 and 2, calculated as the ratio between the length of the proximal region length and the total petal length. Differences were not statistically significant (adjusted P = 0.150 for line #1 and adjusted P = 0.0598 for line #2).

For panels k-n, values in the text are mean ± SD; percentage changes are relative to the wild-type mean. In plots, each point represents one flower-level value, with (n≥9) observations per genotype. All reported (P) values are unadjusted for k. Asterisks indicate significance: **** = p-value < 0.0001). For panels l-n, overall differences were assessed using permutation Kruskal–Wallis tests, with Holm correction. Pairwise comparisons used two-sided exact Wilcoxon–Mann–Whitney tests, with Holm correction across comparisons. Reported pairwise p-values are adjusted.

a b c d

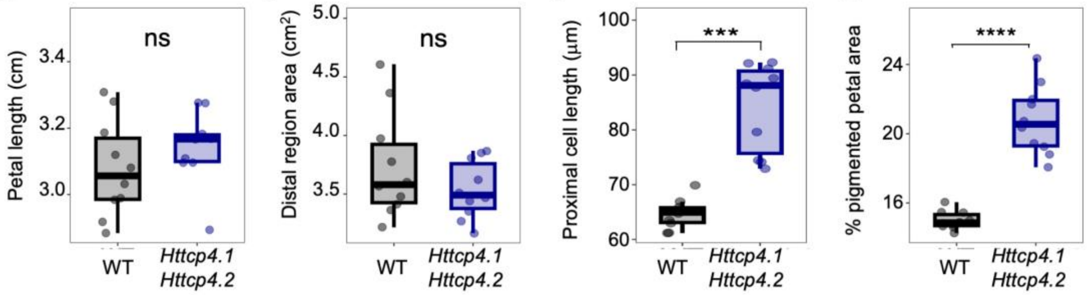


e

| | WT mean | WT SD | *Httcp4.1;Httcp4.2* mean | *Httcp4.1;Httcp4.2* SD | Difference (cell nb) | % Change vs WT |
|---|---|---|---|---|---|---|
| Total cell number | 851 | 21 | 646 | 27 | -205 | -24% |
| Purple cell number | 136 | 8 | 152 | 2 | +16 | ns |
| Cream cell number | 715 | 21 | 494 | 27 | -221 | -31% |
| Tabular cell number | 305 | 19 | 567 | 67 | +262 | +86% |
| Conical cell number | 546 | 27 | 79 | 42 | -467 | -86% |

f

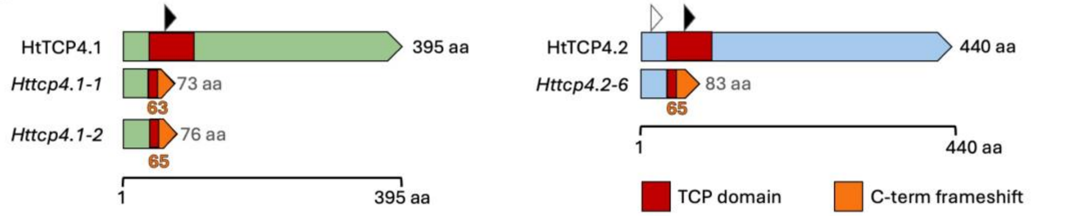


**Supplementary Figure 5.** (**a**) Petal length (cm) in Wild type and *Httcp4.1 Httcp4.2* double-mutant *H. trionum* plants (mean petal length 3.078 ± 0.145 cm vs. 3.143 ± 0.109; adjusted P = 0.800). (**b**) Distal petal area in Wild type and *Httcp4.1 Httcp4.2* double-mutant *H. trionum* plants (mean distal area 3.735 ± 0.453 cm² vs. 3.535 ± 0.246 cm²; Holm-adjusted P= 0.800). (**c**) Length of proximal striated cells in Wild type and *Httcp4.1 Httcp4.2* double-mutant *H. trionum* plants (mean cell length 64.672 ± 2.639 µm vs. 84.233 ± 8.025 µm; adjusted $P = 1.40 \times 10^{-4}$). (**d**) Bullseye proportion, calculated as the proximal-to-total petal area ratio, in Wild type and *Httcp4.1 Httcp4.2* double-mutant *H. trionum* plants (mean bullseye ratio 0.1499 ± 0.0052 vs. 0.2077 ± 0.0199; adjusted $P = 3.94 \times 10^{-5}$). For panel a-d, values are means ± SD; n = 10 per genotype. All comparisons used two-sided Welch's *t*-tests. Boxes indicate the interquartile range, horizontal lines mark medians, and whiskers extend to the most extreme observations within 1.5 times the interquartile range. Wild type (WT) is shown in black and the double mutant (*Httcp4.1 Httcp4.2*) in blue. ns = not significant (p-value ≥ 0.05), *** = p-value < 0.001, **** = p-value < 0.0001. (**e**) Summary of adaxial epidermis cell-count along the proximo-distal petal axis based on cell pigmentation (purple vs cream/white) or cell shape (tabular versus conical) in wild type and *Httcp4.1 Httcp4.2* double mutant. n = 3 per genotype (nb in each sample given on plots in Fig. 5g-h). Values are mean ± SD. Absolute differences and percentage changes are descriptive and relative to wild type. ns = not significant indicate cases where differences between genotypes are not statistically significant, in all other cases p-value <0.0001. (**f**) Schematic representations of the predicted protein truncations in CRISPR/Cas9-edited *HtTCP4.1* and *HtTCP4.2* mutant alleles. Wild-type HtTCP4.1 (395 amino acids) on the left and HtTCP4.2 (440 amino acids) on the right, alongside the predicted truncated proteins encoded by *Httcp4.1-1* (73 amino acids), *Httcp4.1-2* (76 amino acids) and *Httcp4.2-6* (83 amino acids). Red boxes indicate the conserved TCP domain, and orange segments indicate altered amino acid sequences resulting from frameshift mutations. Arrowheads indicate guideRNA target sites (white arrowhead = site targeted by guide1, black arrowhead = site targeted by guide 2). Predicted protein lengths are indicated in grey, start position of altered amino acid sequences due to frameshift mutations are indicated in orange. aa, amino acids. For more details on the mutated genomic sequences, refer to Supplementary Data 5.
